# Alternative developmental and transcriptomic responses to host plant water limitation in a butterfly metapopulation

**DOI:** 10.1101/2021.02.24.432453

**Authors:** Aapo Kahilainen, Vicencio Oostra, Panu Somervuo, Guillaume Minard, Marjo Saastamoinen

## Abstract

Predicting how climate change affects biotic interactions and their evolution poses a challenge. Plant-insect herbivore interactions are particularly sensitive to climate change, as climate-induced changes in plant quality cascade into the performance of insect herbivores. Whereas the immediate survival of herbivore individuals depends on plastic responses to climate change induced nutritional stress, long-term population persistence via evolutionary adaptation requires genetic variation for these responses. In order to assess the prospects for population persistence under climate change, it is therefore crucial to characterise response mechanisms to climate change induced stressors, and quantify their variability in natural populations. Here, we test developmental and transcriptomic responses to water limitation induced host plant quality change in a Glanville fritillary butterfly (*Melitaea cinxia*) metapopulation. We combine nuclear magnetic resonance spectroscopy on the plant metabolome, larval developmental assays and an RNA seq analysis of the larval transcriptome. We observed that responses to feeding on water limited plants, in which amino acids and aromatic compounds are enriched, showed marked intrapopulation variation, with individuals of some families performing better on control and others on water limited plants. The transcriptomic responses were concordant with the developmental responses: Families exhibiting opposite developmental responses also produced opposite transcriptomic responses, e.g. in growth associated intracellular signalling. The opposite developmental and transcriptomic responses are associated with between families differences in organic compound catabolism and storage protein production. The results reveal heritable intrapopulation variability in plasticity, suggesting potential for evolutionary responses to drought-induced changes in host plant quality in the Finnish *M. cinxia* metapopulation.

## Introduction

Because changes in the abiotic environment have different effects on different species, human induced climate change affects species interactions (van Asch & Visser, 2007; Bale et al., 2002; Tylianakis et al., 2008; Voigt et al., 2003). As species interactions are abundant even in the simplest of natural systems (Wirta et al., 2015), much of the effects of climate change are manifested indirectly, rendering any predictions on how climate change affects natural populations difficult (Gilman et al., 2010; Van der Putten et al., 2010; Siepielski et al., 2018; Tylianakis et al., 2008). How much an individuals’ fitness will be affected by quantitative and/or qualitative changes in interacting species depends on their ability to mount plastic responses to compensate for the abrupt changes. In addition, to allow for adaptive evolution at the population level, within population heritable variation in plastic responses is required (Gotthard & Nylin, 1995; Price et al., 2003; Via & Lande, 1985; Wennersten & Forsman, 2012). Therefore, in order to clarify prospects for population persistence under climate change, it is crucial to identify and characterise the response mechanisms associated with changes in interspecific interactions, and examine how they vary within populations.

Interactions between terrestrial insect herbivores and their host plants are particularly susceptible to climate change (van Asch & Visser, 2007; Bale et al., 2002). Climate change not only alters the spatial and phenological availability of host plants, but also modifies their nutrient and secondary metabolite contents (Gershenzon, 1984; Isah, 2019; Rizhsky et al., 2004). These changes in plant quality then cascade to the behaviour and performance of insect herbivores (Jamieson et al., 2012, 2017; Pincebourde et al., 2017) with potential consequences for nutrient cycling and functioning of entire ecosystems (Hunter, 2016; Kalinkat et al., 2015; Post, 2013; Rosenblatt & Schmitz, 2016; Weisser & Siemann, 2008).

Most research on climate change effects on plant-insect herbivore interactions has focused on the effects of changing temperature (Bale et al., 2002; Clissold & Simpson, 2015; Pincebourde et al., 2017). However, more recently, studies on the consequences of changing precipitation have started to accumulate (Jamieson et al., 2012, 2017). Additionally, in agriculture and forestry, there has been a long-lasting interest in understanding the role of host plant water stress in influencing insect pest populations (Cornelissen et al., 2008; Gely et al., 2020; Huberty & Denno, 2004; Larsson, 1989; Mattson & Haack, 1987; Waring & Cobb, 1992; White, 1974). These studies have reported very different responses across systems, with some of the variability attributed to different feeding guilds of the insect herbivores and different drought stress severities across studies.

Nevertheless, considerable variability remains also within the feeding guilds, and very different responses are sometimes found for the same species in different studies (Gutbrodt et al., 2011; Huberty & Denno, 2004; Kuglerová et al., 2019; Rosa et al., 2019; Salgado & Saastamoinen, 2019; Walter et al., 2012). To our knowledge, few studies have examined the contribution of intraspecific and/or intrapopulation variability in insect herbivore responses to host water stress (but see Gibbs et al., 2012). Heritable intrapopulation variability in responses to host plant water stress is also central for the long-term persistence of insect herbivore populations experiencing the effects climate change, as it can allow for evolutionary adaptation to the changing conditions (Carlson et al., 2014; Hoffmann & Sgrò, 2011; Stange et al., 2020; Wennersten & Forsman, 2012).

In order to understand plant-insect herbivore interactions and the persistence of insect herbivore populations during climate change, we need to characterise their response mechanisms to water stress induced changes in plant quality and examine intrapopulation variability therein. To do that, we need to (1) test how water stress changes plant quality, (2) identify insect developmental responses to plant quality, (3) characterise the associated cellular level transcriptomic/genetic responses, and finally (4) quantify how the developmental and transcriptomic responses differ between individuals from different genetic backgrounds (Hoffmann & Sgrò, 2011; Rosenblatt & Schmitz, 2016). Despite long-lasting interest in insect responses to plant water stress (reviewed by Cornelissen, Wilson Fernandes, & Vasconcellos-Neto, 2008; Huberty & Denno, 2004; Larsson, 1989; Waring & Cobb, 1992), and despite the fact that studies on transcriptomic responses of insect herbivores to host plant compounds are emerging (Nallu et al., 2018; Seppey et al., 2019; Vogel et al., 2014), we are not aware of studies combining the two.

Here, we investigate how water stress in the ribwort plantain (*Plantago lanceolata*) cascades into the performance of the Glanville fritillary butterfly (*Melitaea cinxia*) larvae, and how the responses vary within a Finnish metapopulation of the butterfly. Twenty five years of survey data have revealed a change in the metapopulation dynamics of the butterfly, a phenomenon most likely driven by climate change (Hanski & Meyke, 2005; Kahilainen et al., 2018; Tack et al., 2015). Furthermore, recent studies suggest that precipitation across larval stages is positively associated with the regional population growth rate of *M. cinxia* and that drought events can cause abrupt declines in larval population sizes (van Bergen et al., 2020; Kahilainen et al., 2018; Tack et al., 2015).

To examine the mechanisms via which host plant water stress cascades into larval performance, we combined host plant metabolic profiling with development assays and full-transcriptome sequencing of herbivore larvae. First, we profiled metabolic differences between well-watered and water-limited host plants using proton nuclear magnetic resonance spectroscopy (^1^H-NMR). Second, we tested how performance of developing larvae was affected by host plant water limitation. Third, we examined larval gene regulatory responses to water limited host plants by sequencing full transcriptomes of 77 female larvae (RNA seq). Finally, to examine intrapopulation variation in the plastic responses, we compared the phenotypic and transcriptomic responses across full-sib families originating from different parts of the metapopulation.

## Material & Methods

### The ribwort plantain

The ribwort plantain (*Plantago lanceolata, Plantaginaceae*) is the natural host of *M. cinxia* in its studied range. It produces anti-herbivore and anti-fungal chemicals (iridoid glycosides and phenolic compounds) of which the amounts can vary with plant genotype, age, and environmental conditions (Bowers et al., 1992; Bowers & Stamp, 1993).

We collected seeds from a natural population in the Åland Islands (60.196° Lat., 20.704° Lon.) and after germination planted 360 plants in 0.75 litre pots (two saplings each). We reared the plants for three months in controlled greenhouse conditions (ca. 40 ml water/pot daily, 15L:9D photoperiod with 26:18 °C temperature cycle) before initiating the water limitation treatment. We exposed 240 plants to a water limitation treatment in which daily watering was reduced by 50% compared to controls (20ml per pot). This watering scenario was developed in a pilot study in which we experimented with minimum watering allowing the plants to stay alive. More plants were allocated to the water limitation treatment than to the control treatment, because plants produced substantially less leaf biomass in the water limitation treatment (Figure S1). In order to minimize temporal trends in plant quality caused by the plants acclimatising to altered water availability, we initiated the water limitation treatment well in advance (47 days) to the larval exposure (see below).

### Leaf metabolomics assays

Each morning prior to watering (9-10 am), we randomly harvested *P. lanceolata* leaves from control and water limited plants and cut them into 2.25cm^2^ pieces, discarding the basal and tip parts of the leaves. We used these pieces to feed larvae during the experiment (see below), and selected a random subset of six pieces from both treatments for metabolomics assays. For the assays, we recorded the fresh biomass of each piece of leaf, snap froze the pools in liquid nitrogen and stored them in -80 °C. We then freeze dried the samples for 48 hours after which we measured their dry weight, estimated relative water content in the sample [i.e. (fresh mass– dry mass) / fresh mass], and prepared the samples for proton nuclear magnetic resonance spectrometry (^1^H-NMR) following the protocol described by Kim et al. (2010). The ^1^H-NMR spectra of the pool samples were then recorded at the Finnish Biological NMR Center (Institute of Biotechnology, University of Helsinki) and we further processed the obtained spectra for statistical analyses using MNOVA v.10.0.2 software (Mestrelab research S.L., Spain) (Supplementary methods).

### The Glanville fritillary butterfly metapopulation in Finland

The Glanville fritillary butterfly (*Melitaea cinxia*; Lepidoptera: Nymphalidae) is widespread across Eurasia. In Finland, *M. cinxia* is found in the Åland islands archipelago where it exists as a metapopulation inhabiting a network of ca. 4400 discrete habitat patches (mean patch area = 1932 m^2^; sd = 4617 m^2^) containing *P. lanceolata* (Nieminen et al., 2004). The *M. cinxia* metapopulation and its host plants have been surveyed annually since 1993, and the dynamics of the system have been documented in great detail (Hanski et al., 1994, 2017; Nieminen et al., 2004; Ojanen et al., 2013).

In the Åland islands *M. cinxia* is univoltine and the hatching larvae spend their pre-diapause development in gregarious full-sib groups on a single or a couple of host plant individuals and typically enter diapause in the 5^th^ larval instar (Fountain et al., 2017; Kuussaari et al., 2004; Wahlberg, 2000). The pre-diapause larvae are therefore highly susceptible to any changes in host plant quality caused by fluctuations in environmental conditions (Kuussaari et al., 2004).

### Experimental M. cinxia families

We created nine experimental full-sib larval family groups by mating individuals originating from local populations across the natural Åland islands metapopulation (Figure S2). We collected diapausing larvae from nine localities that differed in their average percentages of desiccated host plants (Figure S2, Table S1). We allowed the larvae to continue diapause in a climate chamber (+5 °C, 95% air humidity) for five months and after breaking diapause reared them to adults (12L:12D photoperiod, 28:15 °C temperature cycle) with daily *ad libitum* provision of control reared *P. lanceolata* leaves. To maintain original population genetic structure while minimizing risk of inbreeding in the larval groups, we mated virgin females with males derived from the same habitat patch but different overwintering nests. We placed the mated females on host plants surrounded by mist nets immediately after they had finished mating and collected egg clutches laid on the plant until the female was found dead.

**Table 1.** The association between plant leaf tissue water content, drought treatment and time.

| Covariate | Est. coef. | SE | 95% Cr.I. |  |
| --- | --- | --- | --- | --- |
|  |  |  | Lower | Upper |
| Intercept | 1.033 | 0.053 | 0.931 | 1.137 |
| Water limitation | -0.086 | 0.073 | -0.233 | 0.058 |
| Time | 0.004 | 0.002 | 0.000 | 0.009 |
| Water limitation x Time | 0.001 | 0.003 | -0.005 | 0.008 |

We monitored hatching of egg clutches daily and – in order to have two replicates per full-sib family per treatment – picked two larval groups of a minimum of eighty larvae each from each female to enter the experiment. To minimize potential quality differences caused by clutch rank (Rosa & Saastamoinen, 2017), we selected the larval groups from within the first three larval groups.

### Treatments & developmental assays

On the day after hatching, we divided the larval groups into four smaller groups of twenty larvae each and placed them on separate petri dishes (9 cm diameter, 1.5 cm deep) lined with filter paper. We then randomly assigned each of the dishes to one of four different treatments mimicking different temporal exposures to drought stressed host plants. In addition to a control treatment, in which we fed the larvae with control reared host plants only, the larvae experienced water limited host plant at different stages during pre-diapause development: (1) at late pre-diapause development during 3^rd^ and 4^th^ larval instars, (2) at early pre-diapause development during 1^st^ and 2^nd^ larval instars, and (3) throughout their pre-diapause development from the 1^st^ to the 5^th^ larval instar (Figure S3). In all treatments, we fed the larvae daily with pieces of host plant leaf tissue corresponding to the treatment. We provided *ad libitum* food such that – in order to avoid feeding on old leaf tissue with potentially altered phytochemistry – the larvae consumed most but not all of the leaf tissue during the next 24 hours after provisioning (Supplementary methods).

During the experiment, we recorded development time, body mass at diapause and mortality during development. Once the last larva on a petri dish had entered diapause, we allowed them to spend another four days in normal rearing temperature and photoperiod, after which we measured their body mass and placed them in climate chambers (+5 °C, 95% air humidity) for diapause. We allowed the larvae to diapause for six months, after which we woke them up and recorded overwintering mortality.

With one exception, unexplained mortality during the rearing (i.e. mortality that could not be explained by accidents during handling) was low in all families and in all treatments (mean = 1.1 larvae / petri dish, sd=1.4 larvae). Only the control treatment of the first replicate larval group of family F-5 had a mortality of 55% (11 larvae). As the larvae in this group were developing poorly in general, we concluded it to be an outlier case, potentially suffering from a disease or some other unknown agent, and decided to exclude this larval group from any further analyses.

### Transcriptomics sampling and sequencing

When more than fifty percent of the larvae on a petri dish had spent two full days in the 4^th^ larval instar, we sampled either ten (from the first larval group of each full-sib family) or five (from the second larval group of each full-sib family) larvae for RNA and DNA extraction (see Table S2 for exceptions). Before noon on the day of sampling, we provided the larval groups with leaf tissue matching their treatment and monitored that they fed on the plant before sampling to ensure sampling larvae that are feeding. We weighed and sampled the larvae for RNA and DNA extraction approximately 1.5-2h after feeding, by immersing them in liquid nitrogen and stored the sampled larvae in -80 °C until further processing.

To extract RNA and DNA, we placed the larvae in dry ice and homogenized the frozen larvae in their entirety and separated RNA and DNA following a TRIzol-chloroform purification protocol in combination with QIAamp DNA Mini Kit protocol (Qiagen) (supplementary methods). We then used the extracted DNA for determining the sex of the sampled individuals using sex-specific markers (supplementary methods). To ensure an adequate sample size, we chose to focus only on the transcriptomes of females.

Library preparation from whole RNA and sequencing was conducted at the University Of Helsinki Institute Of Biotechnology (http://www.biocenter.helsinki.fi/bi/). The libraries were prepared using Illumina TruSeq Stranded mRNA Library Prep Kit and sequenced to a depth of a minimum of 13.3 million reads per sample (mean = 17.3M, sd = 1.2M) using Illumina NextSeq 500, with 85bp + 65bp forward and reverse paired-end reads, respectively.

### Sequence data pre-processing, de novo transcriptome assembly, expression quantification

Prior to downstream analyses we removed all Illumina adapter sequence and trimmed low quality sequence using Trimmomatic (version 0.33; Bolger et al. 2014) and verified family structure of the larvae by determining pairwise genetic distances of the individuals from single nucleotide polymorphisms (SNPs) observed in the sequenced reads (supplementary methods).

To build one complete and diverse *de novo* transcriptome, we first built two transcriptomes using the obtained pre-processed reads with both the Trinity (Grabherr et al., 2011; Haas et al., 2013) and Velvet/Oases (Schulz et al., 2012) pipelines. We then combined the two to obtain a single transcriptome of 69 182 putative transcripts using EvidentialGene (Gilbert, 2013). We then mapped the transcriptome against the mitochondrial genome of *M. cinxia* using GMAP (Wu & Watanabe, 2005) and removed all transcripts that exhibited any probability to map to the mitochondrial genome (29 transcripts in total). We then checked the transcriptome for potential contaminants using AAI profiler (Medlar et al., 2018) and electronically annotated the combined transcriptome for transcript protein product descriptions and biological process Gene Ontology terms (BP GO terms; The Gene Ontology Consortium et al., 2000) using PANNZER2 annotation web server (Törönen et al., 2018). For both protein product descriptions and BP GO terms we accepted only annotations above a 0.7 positive predictive value.

Finally, to obtain expected read counts for each sample we mapped the pre-processed reads against the transcriptome using RSEM (B. Li & Dewey, 2011) with Bowtie aligner (Langmead et al., 2009). For further details on building the *de novo* transcriptome and mapping the reads, see supplementary methods.

### Statistical analyses

In order to analyse the proportional water content and the metabolomic response of *P. lanceolata*, we fitted a generalized linear model (GLM) with a Beta-distribution and a constrained correspondence analysis (CCA), respectively. We implemented the GLM following Bayesian inference in the Stan statistical modelling platform (Carpenter et al., 2017) via R (version 3.6.1; R Core Team, 2019) by using packages brms (version 2.7.0; Bürkner, 2017, 2018) and RStan (version 2.17.3; Stan Development Team, 2018). The CCA we implemented using the R package vegan (version 2.5-2; Oksanen et al., 2018). We included the water limitation treatment, days since the start of the experiment, and their interaction with the water limitation treatment as explanatory variables in the GLM and as constraints in the CCA. In addition to the simple temporal trends modelled in the CCA, we tested for convergence of metabolomes in the control and water limited plants. For this, we extracted Euclidean distances between the metabolomes of the two treatments and fitted a Bayesian Gamma distribution GLM in Stan, with days since the beginning of the experiment as an explanatory variable.

We analysed the different phenotypic responses of the larvae by fitting a series of Bayesian generalized linear mixed effects models (GLMMs). We modelled development time using a shifted lognormal distribution, diapause mass using a normal distribution and overwintering mortality using a binomial distribution. All of the models included water limitation treatment, larval family and their interaction as explanatory variables and the egg clutch identity nested within the larval family as a group-level intercept. Like with the above described GLM for water content, we did this in Stan via the R packages brms and Rstan.

Because the majority of larvae in three of the families (F-7, F-8 and F-9; Figure S4, Table S1) entered diapause in the 4^th^ larval instar instead of the 5^th^ (in which we typically observe diapause under laboratory conditions), we analysed the phenotypic responses in families exhibiting primarily 5^th^ instar and 4^th^ instar diapause in separate models. We chose to do this because the distributions of the phenotypic responses of the 4^th^ instar diapausing larvae are widely different from those diapausing in the 5^th^.

Additionally, we observed that the metabolomes of the control and water limited plants converged as the experiment proceeded (see below), with metabolomes being more distinct during the first two instars of the larvae (ca. 10 days; Figure S5). We thus chose to focus on water limitation during early development and combined the control treatment with late water limitation and early water limitation with constant water limitation (Figure S3). Throughout the text we focus primarily on comparisons done for the combined treatments, and refer to these as control and early development water limitation, respectively. We report results differentiating between all temporal water limitation treatments in the supplementary material.

To explore patterns across the transcriptomic dataset, we analysed the effects of family identity, treatment and their interaction on gene expression patterns using redundancy analysis (RDA) as implemented in the R package vegan (version 2.5-2; Oksanen et al., 2018). Prior to model fitting, we normalized the expected count data according to weighted trimmed mean of M-values (TMM; Robinson & Oshlack, 2010) as implemented in the Bioconductor R package edgeR (McCarthy et al., 2012; Robinson et al., 2010), and retained only transcripts with > 1 normalized counts-per-million in a minimum of four samples. Like in the analyses regarding developmental performance, we combined the control treatment with late water limitation and early water limitation with constant water limitation (Figure S3). We then constrained the ordination of the log-transformed counts-per-million values of each transcript with family identity, treatment and their interaction while partialling out potential effects of larval body mass (i.e. model conditional on larval mass).

We approached the biological mechanisms associated with the RDA results in two ways. First, we checked for enrichment of BP GO terms within the set of transcripts among the five percent highest absolute loadings along the RDA axes. Second, we examined individually the annotated protein products in a smaller group of strongest loading percent of the transcripts. For the former approach we checked for enrichment of BP GO using the Bioconductor R package topGO (Alexa & Rahnenfuhrer, 2019) and retained terms using a statistical significance threshold of < 0.01 (Fisher’s exact test with “elim” algorithm; Alexa, Rahnenfuhrer, & Lengauer, 2006). We then clustered the retained BP GO terms hierarchically according to their semantic similarity (Schlicker et al., 2006) using the Bioconductor R package ViSEAGO (Brionne et al., 2019). For illustration of average expression in transcripts closely associated with the enriched BP GO terms, we extracted the average z-score across all transcripts annotated for the enriched GO BP term or its direct offspring terms (including ones not among the 5% with strongest associations). For individual examination of transcripts in the latter approach, we examined transcripts that were annotated for descriptions of protein products and validated the annotations manually with NCBI nucleotide BLAST (Boratyn et al., 2013).

Next, we proceeded to test family-specific transcriptomic responses. For this we fitted transcriptwise negative binomial GLMs with quasi-likelihood F-tests as implemented in the Bioconductor R package edgeR (McCarthy et al., 2012). We conducted filtering and normalization like described above for the RDA and fitted the model such that each family-by-treatment combination was treated as a separate treatment level. Again, to account for the fact that plant metabolites in the different treatments converged during the experiment, we contrasted the coefficients of control and late water limitation with those of early and constant water limitation. We then selected transcripts with a false discovery rate (FDR) below 0.05 and checked for enrichment of BP GO terms using the Bioconductor R package topGO (Alexa & Rahnenfuhrer, 2019) as described above for the RDA across families and treatments.

## Results

### Host plants shift metabolome but not water content upon water limitation treatment

We observed that the leaf tissue water content did not differ between the control and water limited plants (Table 1). However, both control and water limited plants exhibited a tendency for a temporal trend with tissue water content increasing slightly with time. The increase was similar in both treatment levels and differed from zero only once the credible interval was narrowed down to 90% (not shown). The trend suggested a ca. 3% increase in tissue water content during the entire forty day study period.

The metabolic profiles – represented by the sizes of the different ^1^H-NMR chemical shift peaks – differed between the control and water limited plants, and both treatment levels exhibited slightly different temporal trend, as illustrated by the statistically significant treatment (pseudo-F_1,32_=9.759, P<0.001), temporal trend (pseudo-F_1,32_=15.308, P<0.001) and their interaction constraints (pseudo-F_1,32_=3.308, P=0.023) in the CCA model. The CCA model suggested that the two treatments are separated along two statistically significant constrained axes that together account for 45.5% of the variation in the sample metabolite contents (CCA1: pseudo-F_1,32_=23.393, P<0.001; CCA2: pseudo-F_1,32_=3.779, P<0.01; CCA3: pseudo-F_1,32_=1.204, P=0.276; Fig. 2). In both treatment levels the metabolite composition changed in the positive direction along CCA1, whereas the temporal trends were opposite along CCA2 (Figure 2). The overall metabolic profiles of the control and water limited plants converged with time, suggesting that the plants in the two treatments became more similar as the experiment proceeded (Figure S5, Table S3).

A closer examination of the metabolic profiles suggested that water limited plants had higher concentrations of amino acids and aromatic compounds. Signals along the initial (0-3 ppm) and final (6-10 ppm) parts of the ^1^H-NMR spectrum were positively associated with CCA1 (Figure 1b,c). Majority of organic/amino acids and aromatic compounds produce chemical shift peaks within these ranges and, of these, we could identify e.g. glutamine, glutamate, proline, tyrosine and verbascoside (Figure 1b). Several chemical shift peaks within the range typical for carbohydrates were also positively associated with CCA1 but some peaks in this range also had negative associations. Of the positive associations we could identify e.g. xylose, β-D-glucose and catalpol, and of the negative ones sucrose and aucubin (Figure 1b). We could not reliably identify compounds that had strong associations with CCA2, but most of the peaks with strong loadings were within a range where the majority of annotated aromatic compounds and organic acids reside (Figure 1c).

**Figure 1.**
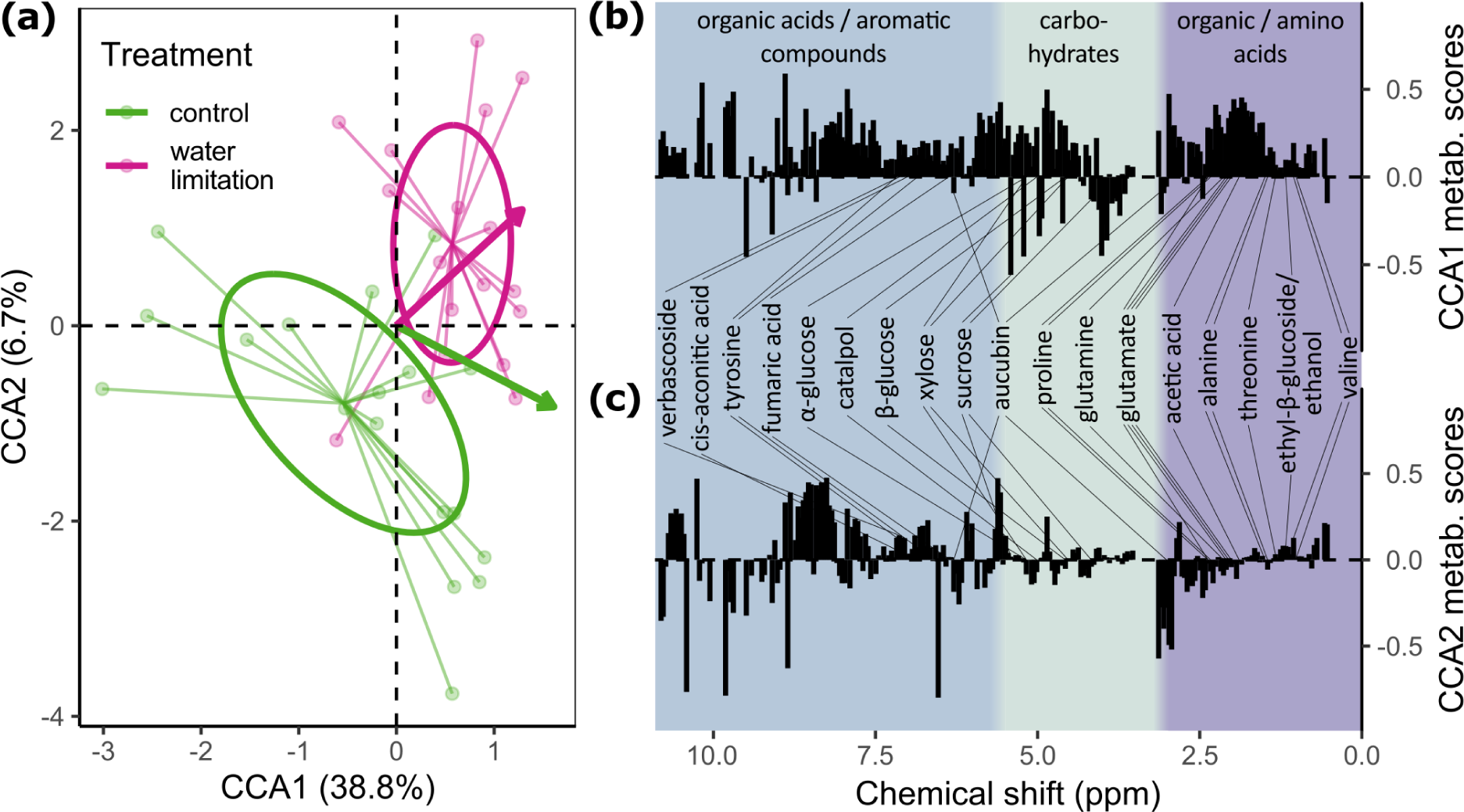
The metabolome differences between control and water limited *P. lanceolata*. (a) The sample scores (points), group centroids (the centre of line spiders) and the 95% confidence interval of the centroids (ellipses) along the first two constrained axes (CCA1 and CCA2) of a constrained correspondence analysis on the associations between plant metabolite composition, water limitation and time since the initiation of the experiment. Arrows depict the temporal trends along the displayed axes. (b,c) The scores of different chemical shift peaks along the first two constrained axes and the compounds the peaks are be associated with. Typical ranges of different types of compounds are shaded with different background colours.

**Figure 2.**
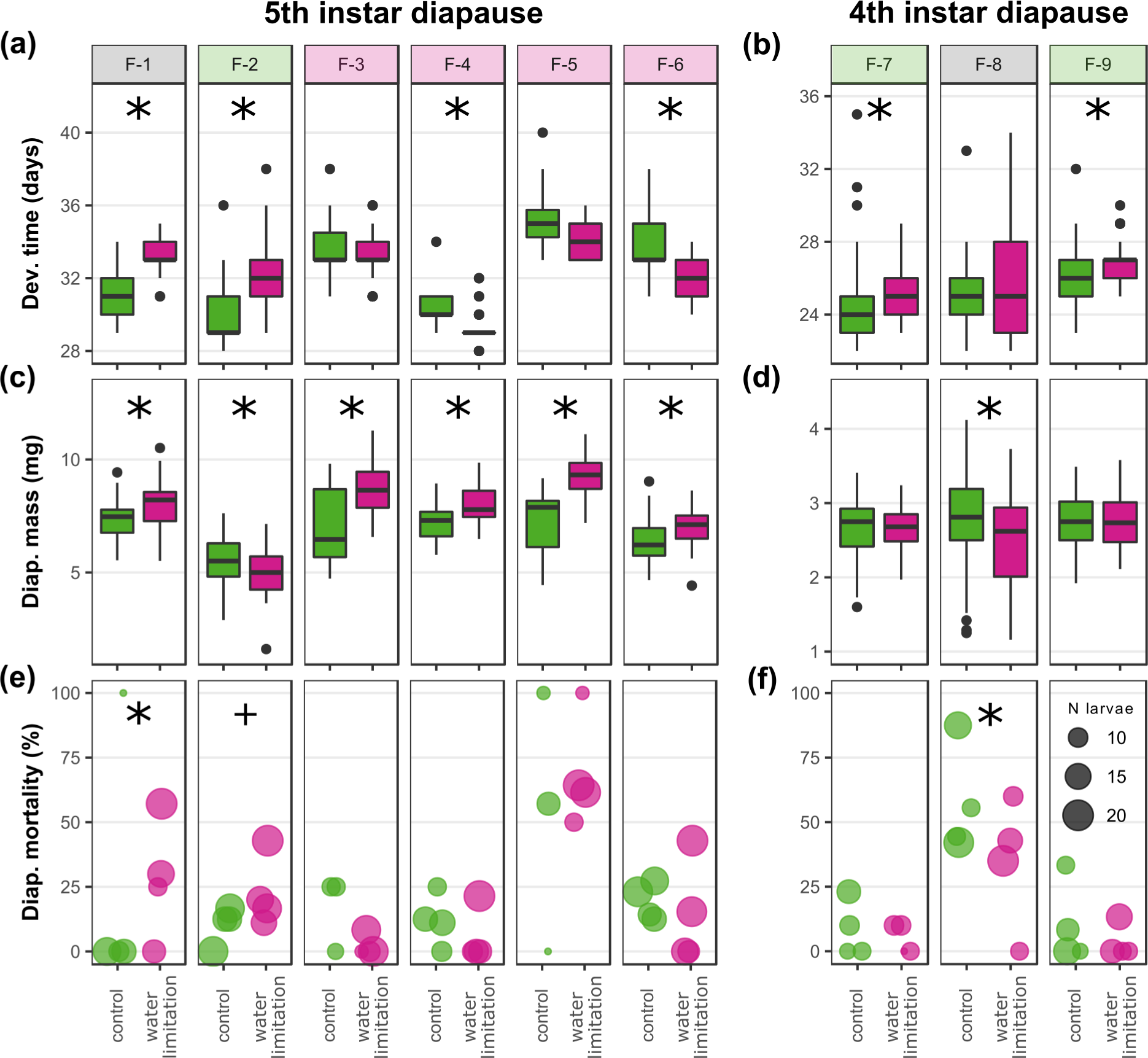
Opposite phenotypic responses of *M. cinxia* to feeding on water limited *P. lanceolata* during the first two instars of their pre-diapause development. Between treatments comparisons in which the 95% and 90% credible intervals of the coefficients do not overlap are indicated with a star and a plus sign, respectively. The panel colour for each family illustrates the treatment in which performance was better. Note the different scales for development time and diapause mass between the 5^th^ (a, c) and 4^th^ instar (b, d) diapause families.

### Variable developmental responses to drought across M. cinxia families

Most of the measured phenotypic traits were influenced by both the family, the host plant water limitation and their interaction, highlighting variability between families in average life history and stress responses (Figure 2, Table S4). We observed three kinds of responses in the studied families: (1) families in which early development water limitation decreased performance in at least one of the phenotypic traits (i.e. increased development time, decreased diapause body mass and / or increased probability of overwintering mortality) (F-2, F-7 and F-9), (2) families in which early development water limitation improved the performance in at least one of the phenotypic traits (F-3, F4, F-5, F-6) and (3) families in which early development water limitation led to mixed responses across traits (F-1 and F-8) (Figure 2). The two families in the last category exhibited larger diapause body masses and higher overwintering mortality when exposed to early development water limitation (Figure 2a,c,d,e,f). Additional analyses maintaining all temporal water limitation levels produced very similar results and supported combining treatment levels according to early water limitation (Figure S6, Table S4).

### Between families transcriptomic differences and divergent transcriptomic responses to water limited host plants

Both the overall transcriptomes and the transcriptomic responses to water limitation differed between *M. cinxia* families. The transcriptomic patterns were parallel to the phenotypic patterns with responses being opposite to each other in some of the families. From the RDA model, we identified family (pseudo-F_3,68_=13.044, P<0.001), treatment (pseudo-F_1,68_=1.703, P=0.033) and their interaction (pseudo-F_3,68_=1.977, P<0.001) as statistically significant constraints, with the constraints together capturing 35.9% of total variance in the data (variance: total = 2385.2, constrained = 856.8, body mass conditioned = 282.6). Of the resulting seven constrained axes, four were statistically significant (RDA1: pseudo-F_1,68_=18.049, P<0.001; RDA2: pseudo-F_1,68_=12.662, P<0.001; RDA3: pseudo-F_1,68_=9.010, P<0.001; RDA4: pseudo-F_1,68_=3.738, P<0.001) and together they explain 37.9% of the variability in the gene expression data (Eigenvalues: RDA1 = 330.7, RDA2 = 232.0, RDA3 = 165.1, RDA4 = 68.5, total = 2102.6).

The first three of these axes illustrate the overall transcriptomic differences between families and the fourth highlights both the responses to host water limitation and the response differences between the families (i.e. a family-by-treatment interaction; Figure 3a, Figure S7). An additional RDA model maintaining all temporal water limitation levels produced practically identical results (Table S6).

**Figure 3.**
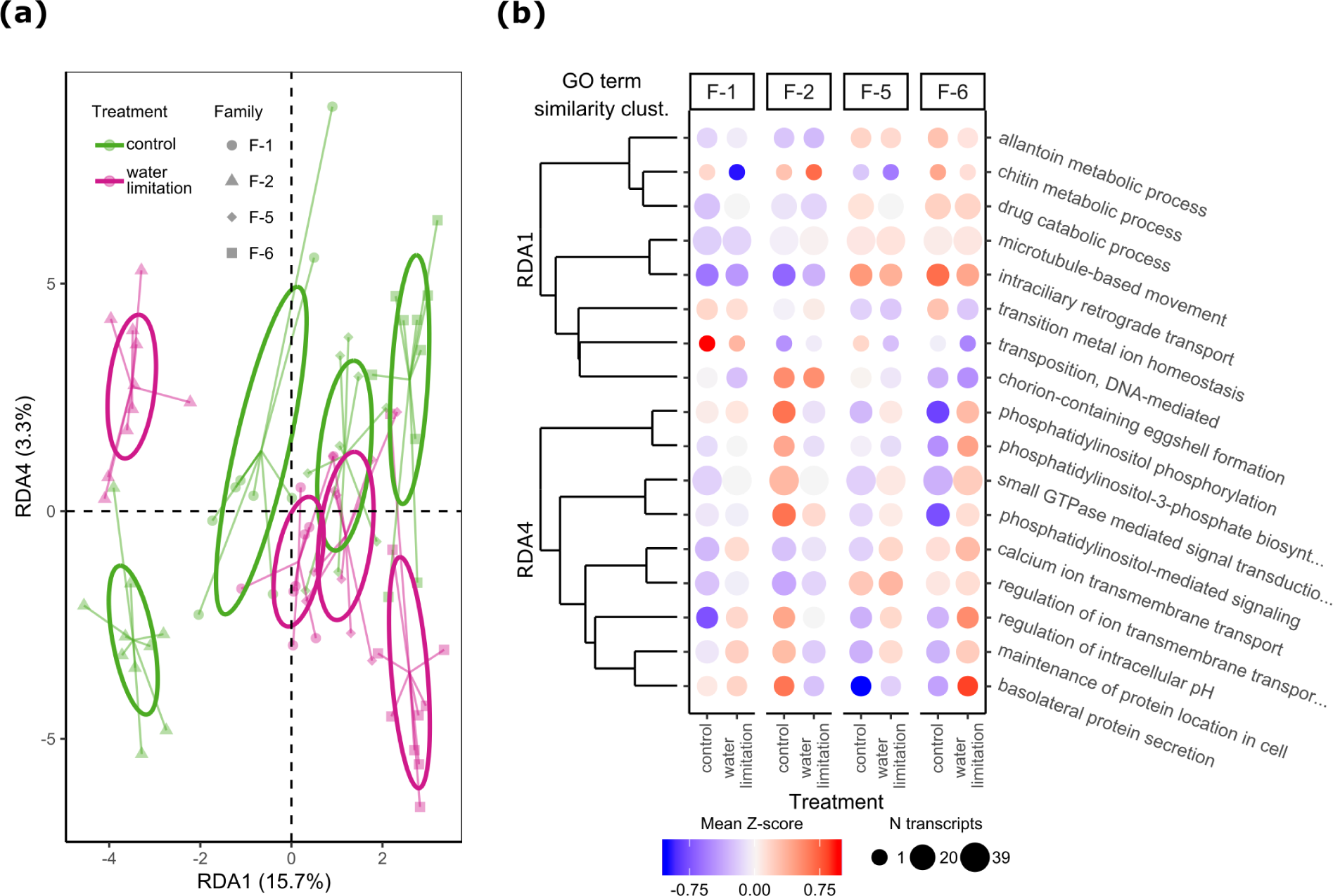
Between families differences in both the overall *M. cinxia* transcriptomic profile and responses to host plant water limitation. (a) The individual scores (points), group centroids (the centre of line spiders) and the 95% confidence interval of the centroids (ellipses) along two constrained axes (RDA1 and RDA4) of an RDA examining the associations between family, treatment and their interaction with the transcriptome of *M. cinxia* larvae. (b) Biological processes enriched in the transcripts showing strongest associations with RDA1 and RDA4, their clustering according to GO-term semantic similarity, and the average expression levels of transcripts associated with the processes. Z-score standardisation of expression levels conducted across families.

The first RDA axis separates families in which performance was improved in the early development water limitation (families F-5 & F-6) from family F-2, in which performance was decreased in the water limitation treatment (Figure 3a). Family F-1, a family that exhibited mixed developmental responses, separates from the others along RDA2 (Figure S7a). For most families, the transcriptomes of the early development water limitation experiencing larvae were negatively associated with RDA4 (and *vice versa* for the control), but for F-2 the pattern was exactly opposite (Figure 3a).

### The transcriptomic differences between families reflect differences in metabolic and growth related processes

Examination of enrichment of BP GO terms within transcripts that contribute most to differences between families (strongest loading 5% of transcripts on RDA1) suggested that families differ in transcripts that contribute to intracellular transportation (i.e. microtubule-based movement and intraciliary retrograde transport), chorion-containing eggshell formation, metabolism of drug-like organic compounds (i.e. drug catabolic process, allantoin metabolic process and chitin metabolic process), transposition, and transition metal ion homeostasis (Figure 3b, Tables S7 and S8). A detailed examination of a smaller set of individual transcripts associated with family differences (strongest loading 1% of transcripts on RDA1) revealed that they produce protein products primarily involved in nutrient storage and development of the sensory system. Transcripts belonging to the former category include those coding for transmembrane protein 135, moderately methionine rich storage protein a, and arylphorin subunit alpha (Table S9). Transcripts belonging to the latter group include those that translate into peripherin-2 like protein, lachesin and gustatory receptor 22 (Table S9). All of the above transcripts are negatively loaded with RDA1 and thus have greatest expression levels in family F-2 and lowest in F-6. The only annotated transcript with a positive loading along RDA1 codes for an intraflagellar transport 52 homolog, which is associated with the assembly, maintenance and function of primary cilia, and is also used in various intracellular functions such as vesicle trafficking and autophagy (Berbari et al., 2009; Finetti et al., 2019; Hua & Ferland, 2018).

The enriched biological processes and protein products associated with RDA4 point towards a stress response that is similar across families, but induced by different treatments in different families. Among the BP GO terms enriched within the strongest loading five percent of transcripts on RDA4, growth associated processes stand out as statistically significant (Figure 3b, Tables S7 and S10). These processes include endocytosis, endosome formation and trafficking, all of which are associated with growth and TOR signalling (Corvera et al., 1999; Flinn et al., 2010; X. Li et al., 2013) (Figure 3b, Table S10). For example, phosphatidylinositol- and small GTPase-mediated signalling play a central role in endocytosis by targeting proteins to the endosome membrane and they are also connected to calcium ion transmembrane transport via inositol phosphates (Berridge, 2016; Corvera et al., 1999; X. Li et al., 2013). Regulation of intracellular pH also points towards endosome related processes, because the acidity of the endosome needs to be adjusted optimal for the functioning of enzymes within the endosome (X. Li et al., 2013). Also, the transcripts that contribute to the enrichment of regulation of intracellular pH are associated with vacuolar acidification, further supporting the conclusion that they contribute to regulation of endosome pH (Table S10).

Detailed examination of individual transcripts (strongest loading 1% of transcripts on RDA4) gives further support to the interpretation that the responses are mostly related to larval growth (Table S11). Some transcripts point towards the production or functioning of specific tissues such as the cuticle (i.e. larval cuticle protein F1-like) or the nervous system (i.e. neurexin-1 alpha) but most transcripts are related to cellular signalling associated with cell proliferation and growth (Table S11). Many protein products belonging to the last category are associated with the serine/threonine-protein kinase TOR pathway (Betz & Hall, 2013; Wullschleger et al., 2006; Zhang et al., 2000). In addition to TOR itself, proteins directly (i.e. headcase protein) and indirectly (i.e. lysine-specific demethylase LID, transient receptor potential cation channel TRPM & inositol 1,4,5-trisphosphate receptor) associated with TOR are coded by the strongest loading 1% of transcripts (Table S11). All the above mentioned transcripts are negatively loaded on RDA4, whereas the opposite is true for transcripts coding for moderately methionine-rich storage protein a, which was also associated with RDA1 (see above). Thus, in family F-2, growth related transcripts are expressed less and moderately methionine-rich storage protein a more when feeding on drought treated plants, and *vice versa* in family F-6 (Figure 3a). In families F-1 and F-5 the responses are qualitatively quite similar to F-6, but less extreme and less consistent between individuals.

### Within families differential gene expression analysis suggests family-specific drought responses

A differential gene expression analysis conducted within each of the families revealed clear family dependency in the transcriptomic response to host plant water limitation (Figure 4). Families showed considerable differences in the numbers and identities of differentially expressed transcripts (Figure 4b). Within most families, individuals grouped quite clearly into the different treatments (Figure 4a). This finding is consistent with the results of the RDA in the sense that individuals in most families exhibit an observable transcriptomic response to early development exposure to drought stressed host (Figure 3a).

**Figure 4.**
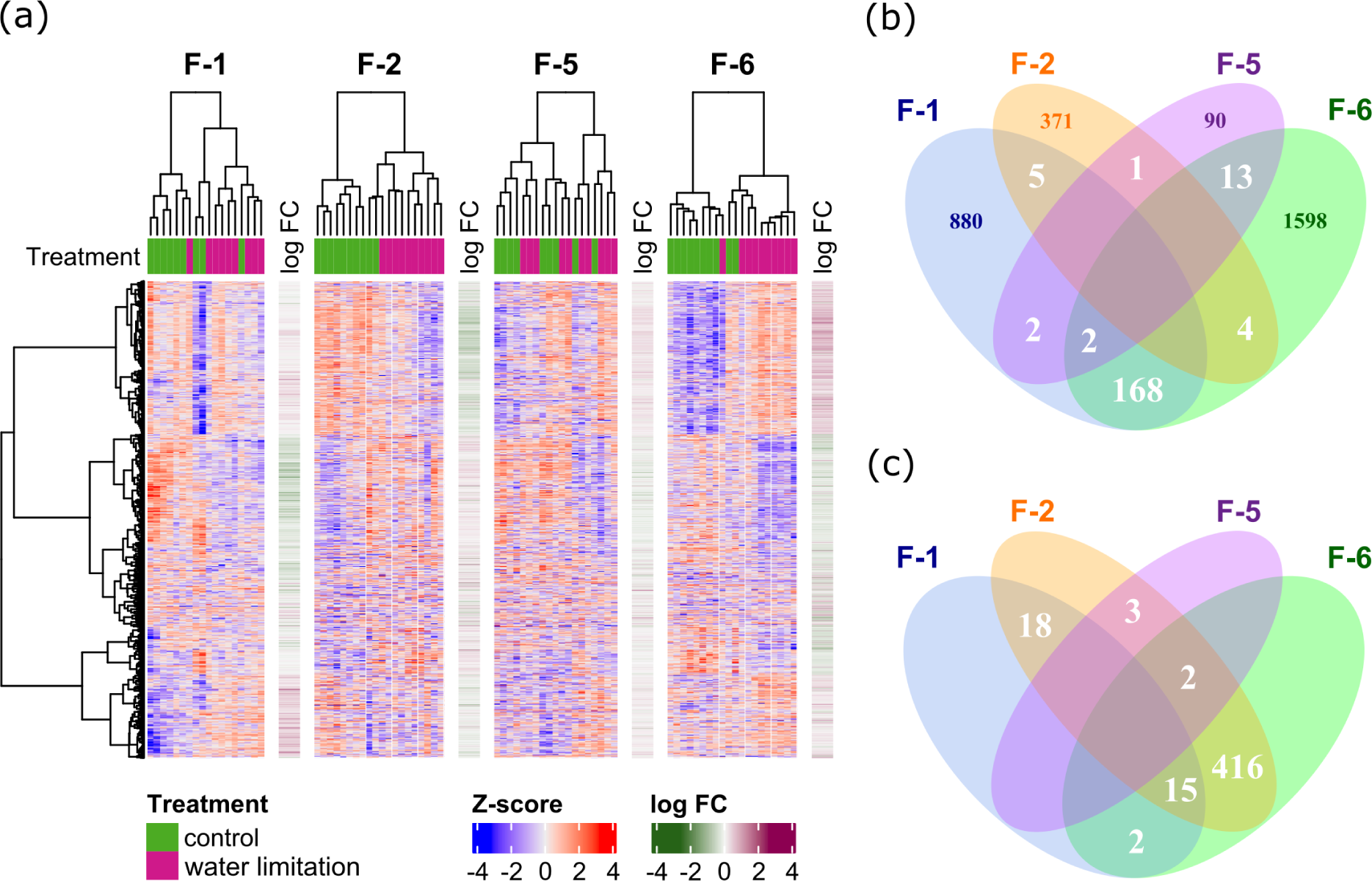
Divergent within-family transcriptomic responses of *M. cinxia* to early development water limitation. (a) A heatmap of differential gene expression in which different panels refer to different larval families with columns and rows representing individuals and transcripts, respectively. The Z-score values in the heatmap (standardised within each family) refer to expression of each transcript in each individual and average expression differences between control and early development water limitation treatment in each transcript in each family are illustrated as log fold-change values (log FC). (b) A Venn diagram for the numbers of transcripts that are differentially expressed in the same direction between control and drought for each family (i.e. either up- or downregulated within the set of families). (c) A Venn diagram for the numbers of differentially expressed transcripts that respond in opposite directions in at least one of the families.

When we focused on differentially expressed transcripts that responded in opposite directions in different families, we notice that family F-2 stands out as responding with many of the same transcripts that are differentially expressed in other families, but the direction of the response is exactly opposite (Figure 4c). In fact, over half of the differentially expressed transcripts in family F-2 are ones that respond in opposite direction in other families. This can also be seen in individual level expression patterns (Figure 4a) and it highlights the family-by-treatment interaction observed along RDA4 (Figure 3a). Thus, both within families differential expression analysis and global RDA identify a transcriptomic stress response that is shared between some of the families but induced by different treatments in different families.

The shared (but opposite) transcriptomic response between families F-2 and F-6 is also reflected in enrichment of similar BP GO-terms (Figure 5, Table S12). Concordant with the RDA, intracellular signalling related to endocytosis and endosome trafficking (small GTPase- and phosphatidylinositol-mediated signalling) are prominently influenced by feeding on water limited host plant in these families. Furthermore, the fact that both families also respond with protein (de-)ubiquitination related transcripts, further strengthens the interpretation of involvement of endocytosis and endosome trafficking, because ubiquitin can serve as a targeting que in endosome trafficking (Piper & Lehner, 2011) (Figure 5, Table S12).

**Figure 5.**
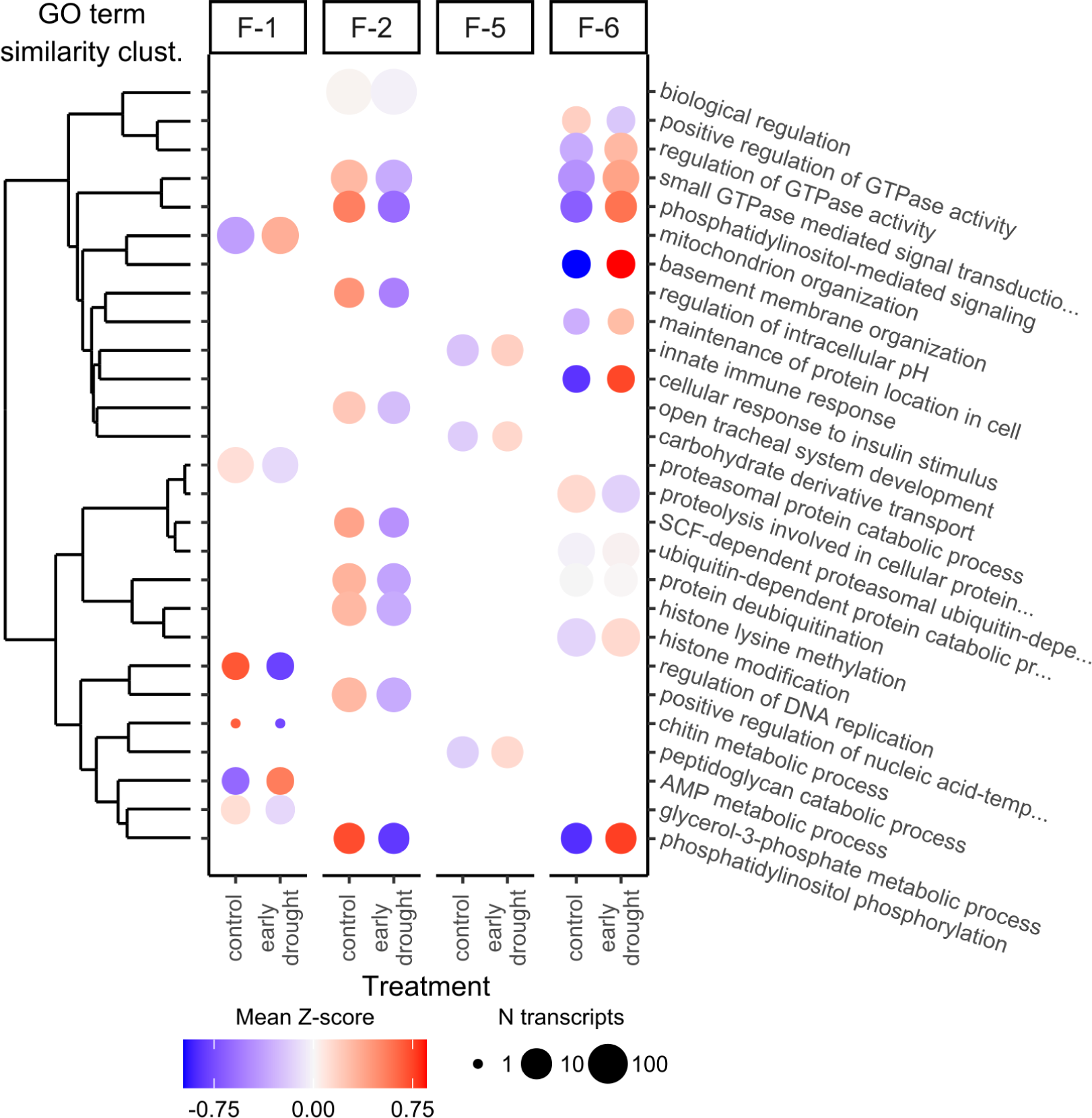
Biological processes associated with divergent within-family responses of *M. cinxia* to early development water limitation. For each family, only the statistically significantly (P<0.01) enriched BP GO terms are included. Z-score standardisation of expression levels conducted within families.

In families F-1 and F-5, the processes were not as clearly associated with endocytosis. Nevertheless, the BP GO-terms enriched in family F-1 (i.e. regulation of DNA replication, proteasomal protein catabolism, and metabolism of chitin, AMP and glycerol-3-phosphate; Figure 5) suggest differences in cell replication and in nutrient metabolism between the control and water limited host plant feeding individuals. The few enriched processes in F-5 are associated with immune responses (Figure 5), which could indicate a general stress response or exposure to infections. Based on the developmental responses, and high pre-diapause mortality in the control treatment of another larval group replicate of the same family, one would expect stress/infection related responses in individuals feeding on the control treated plants (Figure 2). The expression patterns, however, do not support this as the immunity related responses seems to be upregulated in the water limitation treatment (Figure 5), in which the developmental performance of this family was improved.

## Discussion

We discovered that the Finnish *Melitaea cinxia* metapopulation harbours divergent plastic responses to *Plantago lanceolata* nutritional quality induced by changes in water limitation. First, we found that in some full-sib families larval performance was improved when feeding on water limited plants during early larval developmental stages, whereas the exact opposite was true for other families. Second, we observed transcriptomic patterns parallel to the divergent developmental responses revealing a stress response in which growth related signalling processes were downregulated similarly across most studied families. However, in some families the response was induced by the water limited plants whereas in others by control treated plants. Third, we observed between families transcriptomic differences associated with the divergent phenotypic responses and – consistent with observed water limitation induced changes in plant quality – discovered that they reflect differences in nutrient storage, metabolising organic compounds and intracellular transport.

### Intrapopulation variability in performance on water stressed host plants

We found that whereas the larvae of four *M. cinxia* families grew larger, developed faster and/or had lower overwintering mortality when feeding on water stressed host plants in which amino acids and phenolic compounds were enriched, the opposite was true for three of the families, and mixed responses were observed in two families (Figures 1 and 2). The observation of opposite response types in approximately equal proportions in a single metapopulation suggests that intraspecific variability in insect herbivore responses to host plant water stress can be considerable in natural systems.

Although abundance of studies have identified several factors influencing insect herbivores’ responses to host plant water stress, including the feeding guild of the insect, the severity and duration of water stress, and phenology of the system (Che-Castaldo et al., 2019; Cornelissen et al., 2008; Huberty & Denno, 2004; Larsson, 1989; Waring & Cobb, 1992; White, 2009; Yang et al., 2020), our results are among the first to highlight the importance of intraspecific variability in insect water stress responses (but see Gibbs et al., 2012). Such intraspecific variability might explain some of the contradicting results reported for the same species in different studies. In fact, two of our recent studies utilizing very similar experimental designs have reported opposite responses of the Finnish *M. cinxia* to water stressed *P. lanceolata* (Rosa et al., 2019; Salgado & Saastamoinen, 2019). However, the design in these studies was not aimed at capturing variability across the range of the metapopulation, and thus it is possible that they sampled families with different developmental strategies by chance. Similar between studies discrepancies have been reported also for Spodoptera littoralis, but here comparisons are more difficult as the experimental manipulations and host plants vary between studies (Gutbrodt et al., 2011; Kuglerová et al., 2019; Walter et al., 2012).

### Parallel transcriptomic and developmental responses reveal a stress response to sub-optimal nutrition

Full transcriptome sequencing (RNA seq) of 77 female larvae belonging to four of the nine studied families revealed a transcriptomic response parallel to the divergent phenotypic responses. A redundancy analysis (RDA) highlighted divergent transcriptomic responses to water limited plants between the larval families performing better on control and water limited plants (Figure 3a). A similar divergent pattern was also observed in a transcriptwise differential gene expression analysis conducted within each of the families (Figure 4a,c).

Although *M. cinxia* larval families exhibited opposite transcriptomic responses to feeding on water limited plants, the fact that individuals exhibiting reduced developmental performance responded quite similarly with several transcripts points towards a stress response that is shared among families (Figures 2-4). This view is strengthened by the fact that the transcripts associated with the divergent responses were involved in growth related processes, with larvae exhibiting reduced developmental performance having lower expression of transcripts coding for intracellular signalling proteins involved in endocytosis and the serine/theroninen-protein kinase TOR (TOR) pathway (Figures 3b and 5, Tables S7, S10-S12) (Corvera et al., 1999; Flinn et al., 2010; X. Li et al., 2013).

The TOR pathway is associated with increased cell proliferation and growth during good nutrient conditions (Betz & Hall, 2013; N. Li et al., 2019; Scott et al., 2004) and suppression of genes associated with TOR signalling indicates a stress response to sub-optimal nutrient environment. Indeed, a recent study discovered that inhibiting the expression of TOR slowed down the growth rate of the larvae of the lepidopteran *Maruca virata* and further verified that the expression of TOR varied in concert with growth rate on different alternative host plants (Al Baki et al., 2018).

### Between families transcriptomic differences underlie divergent responses to host plant water stress

Between families transcriptomic differences provides candidate processes and pathways underlying the divergent developmental and transcriptomic responses to host plant water limitation. In the RDA across the RNA seq dataset most of the variability (15.7%) was explained by differences between families (Figure 3a). Notably, the first axis of the RDA separated families exhibiting opposite phenotypic responses to opposite ends of the axis (Figures 2 and 3a, Table S4).

The protein products associated with the between families differences (i.e. those most strongly loaded on the first RDA axis) revealed differences regarding metabolising and storing nutrients. Individuals of the family F-2, in which performance was better on control treated plants, had higher baseline expression of transcripts involved in the production of storage proteins (moderately methionine-rich protein a and arylphorin) and fat accumulation (transmembrane protein 135) (Table S9). In holometabolous insects, these proteins are typically associated with preparing for metamorphosis, diapause, egg production, or periods of environmental/nutritional stress (Ashfaq et al., 2007; Denlinger, 2000; Exil et al., 2010; Haunerland, 1996; Sonoda et al., 2007). It is thus possible that individuals in family F-2 allocated more resources to nutrient storage than individuals in other families. The diapause body masses of individuals in family F-2 were generally lower than those of individuals in other families, which may be an indication that the increased nutrient storage may have come with an expense on growth (Figure 2c, Table S4). Alternatively, individuals of the family F-2 may have experienced the conditions generally more stressful, which led to smaller diapause body mass and increased storage protein production.

In families F-5 and F-6, individuals benefitted from feeding on water limited plants. They had higher average expression of transcripts associated with metabolizing drug-like compounds and particularly catabolism of allantoin, a purine metabolism product (Figure 3, Table S7) (Bursell, 1967). Increased allantoin catabolism may be an indication of more efficient breakdown of amino acids allowing for better nitrogen intake when feeding on the amino acid rich water limited plants (Figure 1). Also, in several plant species, allantoin is known to become enriched in plant tissues during water limitation and better allantoin catabolism may have directly enabled increased nitrogen availability when feeding on drought stressed hosts (Bowne et al., 2012; Casartelli et al., 2019; Coleto et al., 2014; Oliver et al., 2011). Unfortunately, we cannot explicitly single out accumulation of allantoin in our drought stressed plants due to overlapping peaks in its ^1^H-NMR chemical shift range.

Families F-5 and F-6 also had higher average expression of transcripts associated with intraciliary and microtubule-based movement (Figure 3, Table S7). This could be related to the observed allantoin catabolism and purine metabolism in general, because purine catabolism typically occurs in the peroxisome (Islinger et al., 2010), which are transported along microtubules (Hancock, 2014). Furthermore, purinosomes – enzyme clusters involved in purine synthesis – have recently been shown to be tightly associated with microtubule-based movement (Chan et al., 2018). However, the connection between higher baseline allantoin catabolism and intracellular transportation in families benefitting from water limited plants should be interpreted with caution, because microtubule-based movement and intraflagellar transport (IFT) system are involved in a wide range of cellular processes such as cell cycle control, endocytosis, autophagy and vesicle trafficking (Berbari et al., 2009; Finetti et al., 2019; Hancock, 2014; Hua & Ferland, 2018).

### A butterfly in a changing world

Like so many natural systems, the Åland islands *M. cinxia* metapopulation is currently threatened by human induced climate change. In a long-term time series of over 27 years of demographic records, the last 15 years have marked a change in the metapopulation dynamics of the system, with increasing fluctuations in abundance elevating the extinction risk (van Bergen et al., 2020; Hanski & Meyke, 2005; Kahilainen et al., 2018; Tack et al., 2015). Although we know that regional population growth rates are linked with precipitation across larval stages (van Bergen et al., 2020; Kahilainen et al., 2018; Tack et al., 2015), a detailed understanding of individual-level responses to water limitation (and variability therein) is required in order to predict the behaviour of the system and design necessary conservation actions.

The observed between families variability in responses to host plant water limitation is a promising sign of heritable genetic variability in drought tolerance within the system. However, since the parental generation of the current study was collected from the wild, we cannot completely rule out the possibility that transgenerational effects might be influencing the observed divergent transcriptomic and developmental responses. Our previous work suggests that adult condition is primarily influenced by environmental conditions during post-diapause development (Saastamoinen et al., 2013) and by allowing the parental generation to spend their diapause period and post-diapause development in controlled laboratory conditions we have aimed to minimize potential transgenerational effects induced by quality differences in the parental generation.

The Åland islands *M. cinxia* metapopulation has been reported to contain heritable variation in behavioural and developmental responses to different environmental variables (e.g. temperature) across different life-history stages (Kvist et al., 2013; Niitepõld & Saastamoinen, 2017; Verspagen et al., 2020). It has been suggested that metapopulation dynamics, with frequent colonisations and extinctions, maintain genetic variability in plastic responses via complex trade-offs between dispersal ability and other life-history characteristics (Kvist et al., 2013; Niitepõld & Saastamoinen, 2017). Although not explicitly the focus of the current study, it is possible that similar trade-offs are also associated with the observed variability in responses to drought stressed host plants.

## Conclusions

We observe very different including opposite performance and transcriptomic responses across different *M. cinxia* larval full-sib families. This suggests that intraspecific variability can contribute to the variety of observed responses of plant-herbivore systems to changing water availability in previous studies. The observed inter-family variation provides evidence for variation in phenotypic plasticity within the Finnish *M. cinxia* metapopulation, potentially improving its chances of persisting in the changing climate. Our results highlight the importance of unravelling the magnitude and genetic mechanisms behind intrapopulation variability for understanding and predicting the abilities of natural populations to respond to climate change.

## Acknowledgements

We thank Suvi Ikonen, Auli Relve, Heini Karvinen and Alma Oksanen for helping with the practical work related to rearing the larvae, and Laura Häkkinen and Juha-Matti Pitkänen for helping with sample preparations. In addition, we thank Virpi Ahola, Mikko Frilander and Jouni Kvist for valuable comments when designing the study. Transcriptomic analyses were conducted with the aid of computational resources provided by CSC – IT Center for Science, Finland. The NMR facility at the Institute of Biotechnology, University of Helsinki is supported by Biocenter Finland and Helsinki Institute of Life Science (HiLIFE). The study was funded by European Research Council (Independent starting grant no. 637412 ‘META-STRESS’ to MS) and Kone Foundation (grant no. 201802795 to AK).

## Data Accessibility

RNA seq reads, raw and processed read count tables and transcriptome assembly are available from NCBI’s Gene Expression Omnibus, with the accession number GSE159376.

## Author Contributions

AK & MS developed the study questions and study design, AK conducted the experiment, AK, VO, PS & GM analysed the data, AK wrote the first version of the manuscript to which all authors contributed.

## Supplementary material

Supplementary figures and tables

Supplementary methods

## References

Alexa, A., & Rahnenfuhrer, J. (2019). topGO: enrichment analysis for Gene Ontology. Retrieved from https://bioconductor.org/packages/release/bioc/html/topGO.html

Alexa, A., Rahnenfuhrer, J., & Lengauer, T. (2006). Improved scoring of functional groups from gene expression data by decorrelating GO graph structure. Bioinformatics, 22(13), 1600–1607. https://doi.org/10.1093/bioinformatics/btl140

van Asch, M., & Visser, M. E. (2007). Phenology of forest caterpillars and their host trees: The importance of synchrony. Annual Review of Entomology, 52(1), 37–55. https://doi.org/10.1146/annurev.ento.52.110405.091418

Ashfaq, M., Sonoda, S., & Tsumuki, H. (2007). Expression of two methionine-rich storage protein genes of Plutella xylostella (L.) in response to development, juvenile hormone-analog and pyrethroid. Comparative Biochemistry and Physiology Part B: Biochemistry and Molecular Biology, 148(1), 84–92. https://doi.org/10.1016/J.CBPB.2007.04.017

Al Baki, M. A., Jung, J. K., Maharjan, R., Yi, H., Ahn, J. J., Gu, X., & Kim, Y. (2018). Application of insulin signaling to predict insect growth rate in Maruca vitrata (Lepidoptera: Crambidae). PLo ONE, 13(10), e0204935. https://doi.org/10.1371/journal.pone.0204935

Bale, J. S., Masters, G. J., Hodkinson, I. D., Awmack, C., Bezemer, T. M., Brown, V. K., et al. (2002). Herbivory in global climate change research: direct effects of rising temperature on insect herbivores. Global Change Biology, 8(1), 1–16. https://doi.org/10.1046/j.1365-2486.2002.00451.x

Berbari, N. F., O’Connor, A. K., Haycraft, C. J., & Yoder, B. K. (2009). The primary cilium as a complex signaling center. Current Biology, 19(13), R526–R535. https://doi.org/10.1016/j.cub.2009.05.025

van Bergen, E., Dallas, T., DiLeo, M. F., Kahilainen, A., Mattila, A. L. K., Luoto, M., & Saastamoinen, M. (2020). The effect of summer drought on the predictability of local extinctions in a butterfly metapopulation. Conservation Biology, 34(6), 1503–1511. https://doi.org/10.1111/cobi.13515

Berridge, M. J. (2016). The inositol trisphosphate/calcium signaling pathway in health and disease. Physiological Reviews, 96(4), 1261–1296. https://doi.org/10.1152/physrev.00006.2016

Betz, C., & Hall, M. N. (2013). Where is mTOR and what is it doing there? Journal of Cell Biology, 203(4), 563–574. https://doi.org/10.1083/jcb.201306041

Bolger, A. M., Lohse, M., & Usadel, B. (2014). Trimmomatic: a flexible trimmer for Illumina sequence data. Bioinformatics, 30(15), 2114–2120. https://doi.org/10.1093/bioinformatics/btu170

Boratyn, G. M., Camacho, C., Cooper, P. S., Coulouris, G., Fong, A., Ma, N., et al. (2013). BLAST: a more efficient report with usability improvements. Nucleic Acids Research, 41(W1), W29–W33. https://doi.org/10.1093/nar/gkt282

Bowers, M. D., & Stamp, N. E. (1993). Effects of plant age, genotype and herbivory on Plantago performance and chemistry. Ecology, 74(6), 1778–1791. https://doi.org/10.2307/1939936

Bowers, M. D., Collinge, S. K., Gamble, S. E., & Schmitt, J. (1992). Effects of genotype, habitat, and seasonal variation on iridoid glycoside content of Plantago lanceolata (Plantaginaceae) and the implications for insect herbivores. Oecologia, 91(2), 201–207. https://doi.org/10.1007/BF00317784

Bowne, J. B., Erwin, T. A., Juttner, J., Schnurbusch, T., Langridge, P., Bacic, A., & Roessner, U. (2012). Drought responses of leaf tissues from wheat cultivars of differing drought tolerance at the metabolite level. Molecular Plant, 5(2), 418–429. https://doi.org/10.1093/MP/SSR114

Brionne, A., Juanchich, A., & Hennequet-Antier, C. (2019). ViSEAGO: a Bioconductor package for clustering biological functions using Gene Ontology and semantic similarity. BioData Mining, 12(1), 16. https://doi.org/10.1186/s13040-019-0204-1

Bürkner, P.-C. (2017). brms: An R Package for Bayesian Multilevel Models using Stan. Journal of Statistical Software, 80(1), 1–28. https://doi.org/10.18637/jss.v080.i01

Bürkner, P.-C. (2018). Advance Bayesian Multilevel Modeling with the R Package brms. The R Journal, 10(1), 395–411.

Bursell, E. (1967). The excretion of nitrogen in insects. Advances in Insect Physiology, 4, 33– 67. https://doi.org/10.1016/S0065-2806(08)60207-6

Carlson, S. M., Cunningham, C. J., & Westley, P. A. H. (2014). Evolutionary rescue in a changing world. Trends in Ecology and Evolution, 29(9), 521–530. https://doi.org/10.1016/j.tree.2014.06.005

Carpenter, B., Gelman, A., Hoffman, M. D., Lee, D., Goodrich, B., Betancourt, M., et al. (2017). Stan: a probabilistic programming language. Journal of Statistical Software, 76(1), 1–32. https://doi.org/10.18637/jss.v076.i01

Casartelli, A., Melino, V. J., Baumann, U., Riboni, M., Suchecki, R., Jayasinghe, N. S., et al. (2019). Opposite fates of the purine metabolite allantoin under water and nitrogen limitations in bread wheat. Plant Molecular Biology, 99(4–5), 477–497. https://doi.org/10.1007/s11103-019-00831-z

Chan, C. Y., Pedley, A. M., Kim, D., Xia, C., Zhuang, X., & Benkovic, S. J. (2018). Microtubule-directed transport of purine metabolons drives their cytosolic transit to mitochondria. Proceedings of the National Academy of Sciences of the United States of America, 115(51), 13009–13014. https://doi.org/10.1073/pnas.1814042115

Che-Castaldo, C., Crisafulli, C. M., Bishop, J. G., Zipkin, E. F., & Fagan, W. F. (2019). Disentangling herbivore impacts in primary succession by refocusing the plant stress and vigor hypotheses on phenology. Ecological Monographs, 89(4), e01389. https://doi.org/10.1002/ecm.1389

Clissold, F. J., & Simpson, S. J. (2015). Temperature, food quality and life history traits of herbivorous insects. Current Opinion in Insect Science, 11, 63–70. https://doi.org/10.1016/j.cois.2015.10.011

Coleto, I., Pineda, M., Rodiño, A. P., De Ron, A. M., & Alamillo, J. M. (2014). Comparison of inhibition of N2 fixation and ureide accumulation under water deficit in four common bean genotypes of contrasting drought tolerance. Annals of Botany, 113(6), 1071–1082. https://doi.org/10.1093/aob/mcu029

Cornelissen, T., Wilson Fernandes, G., & Vasconcellos-Neto, J. (2008). Size does matter: variation in herbivory between and within plants and the plant vigor hypothesis. Oikos, 117(8), 1121–1130. https://doi.org/10.1111/j.0030-1299.2008.16588.x

Corvera, S., D’Arrigo, A., & Stenmark, H. (1999). Phosphoinositides in membrane traffic. Current Opinion in Cell Biology, 11(4), 460–465. https://doi.org/10.1016/S0955-0674(99)80066-0

Denlinger, D. L. (2000). Molecular Regulation of Insect Diapause. In K. B. Storey & J. M. Storey (Eds.), Cell and Molecular Response to Stress (pp. 259–275). Elsevier. https://doi.org/10.1016/S1568-1254(00)80020-0

Exil, V. J., Silva Avila, D., Benedetto, A., Exil, E. A., Adams, M. R., Au, C., & Aschner, M. (2010). Stressed-induced TMEM135 protein is part of a conserved genetic network involved in fat storage and longevity regulation in Caenorhabditis elegans. PLoS ONE, 5(12), e14228. https://doi.org/10.1371/journal.pone.0014228

Finetti, F., Capitani, N., & Baldari, C. T. (2019). Emerging roles of the intraflagellar transport system in the orchestration of cellular degradation pathways. Frontiers in Cell and Developmental Biology, 7, 292. https://doi.org/10.3389/fcell.2019.00292

Flinn, R. J., Yan, Y., Goswami, S., Parker, P. J., & Backer, J. M. (2010). The late endosome is essential for mTORC1 signaling. Molecular Biology of the Cell, 21(5), 833–841. https://doi.org/10.1091/mbc.E09-09-0756

Fountain, T., Husby, A., Nonaka, E., DiLeo, M. F., Korhonen, J. H., Rastas, P., et al. (2017). Inferring dispersal across a fragmented landscape using reconstructed families in the Glanville fritillary butterfly. Evolutionary Applications, 11(3), 287–297. https://doi.org/10.1111/eva.12552

Gely, C., Laurance, S. G. W., & Stork, N. E. (2020). How do herbivorous insects respond to drought stress in trees? Biological Reviews, 95(2), 434–448. https://doi.org/10.1111/brv.12571

Gershenzon, J. (1984). Changes in the levels of plant secondary metabolites under water and nutrient stress. In B. N. Timmermann, C. Steelink, & F. A. Loewus (Eds.), Phytochemical Adaptations to Stress (pp. 273–320). Boston, MA: Springer US. https://doi.org/10.1007/978-1-4684-1206-2_10

Gibbs, M., Van Dyck, H., & Breuker, C. J. (2012). Development on drought-stressed host plants affects life history, flight morphology and reproductive output relative to landscape structure. Evolutionary Applications, 5(1), 66–75. https://doi.org/10.1111/j.1752-4571.2011.00209.x

Gilbert, D. (2013, June). EvidentialGene. 7th Annual Arthropod Genomics Symposium. Notre Dame: Indiana University. https://doi.org/10.7490/F1000RESEARCH.1112594.1

Gilman, S. E., Urban, M. C., Tewksbury, J., Gilchrist, G. W., & Holt, R. D. (2010). A framework for community interactions under climate change. Trends in Ecology and Evolution, 25(6), 325–331. https://doi.org/10.1016/j.tree.2010.03.002

Gotthard, K., & Nylin, S. (1995). Adaptive plasticity and plasticity as an adaptation: A selective review of plasticity in animal morphology and life history. Oikos, 74(1), 3–17. https://doi.org/10.2307/3545669

Grabherr, M. G., Haas, B. J., Yassour, M., Levin, J. Z., Thompson, D. A., Amit, I., et al. (2011). Full-length transcriptome assembly from RNA-Seq data without a reference genome. Nature Biotechnology, 29(7), 644–652. https://doi.org/10.1038/nbt.1883

Gutbrodt, B., Mody, K., & Dorn, S. (2011). Drought changes plant chemistry and causes contrasting responses in lepidopteran herbivores. Oikos, 120(11), 1732–1740. https://doi.org/10.1111/j.1600-0706.2011.19558.x

Haas, B. J., Papanicolaou, A., Yassour, M., Grabherr, M., Blood, P. D., Bowden, J., et al. (2013). *de novo* transcript sequence reconstruction from RNA-seq using the Trinity platform for reference generation and analysis. Nature Protocols, 8(8), 1494–1512. https://doi.org/10.1038/nprot.2013.084

Hancock, W. O. (2014). Bidirectional cargo transport: Moving beyond tug of war. Nature Reviews Molecular Cell Biology, 15(9), 615–628. https://doi.org/10.1038/nrm3853

Hanski, I., & Meyke, E. (2005). Large-scale dynamics of the Glanville fritillary butterfly: Landscape structure, population processes, and weather. Annales Zoologici Fennici, 42, 379–395.

Hanski, I., Kuussaari, M., & Nieminen, M. (1994). Metapopulation structure and migration in the butterfly Melitaea cinxia. Ecology, 75(3), 747–762. https://doi.org/10.2307/1941732

Hanski, I., Schulz, T., Wong, S. C., Ahola, V., Ruokolainen, A., & Ojanen, S. P. (2017). Ecological and genetic basis of metapopulation persistence of the Glanville fritillary butterfly in fragmented landscapes. Nature Communications, 8, 14504. https://doi.org/10.1038/ncomms14504

Haunerland, N. H. (1996). Insect storage proteins: Gene families and receptors. Insect Biochemistry and Molecular Biology, 26(8–9), 755–765. https://doi.org/10.1016/S0965-1748(96)00035-5

Hoffmann, A. A., & Sgrò, C. M. (2011). Climate change and evolutionary adaptation. Nature, 470(7335), 479–485. https://doi.org/10.1038/nature09670

Hua, K., & Ferland, R. J. (2018). Primary cilia proteins: ciliary and extraciliary sites and functions. Cellular and Molecular Life Sciences, 75(9), 1521–1540. https://doi.org/10.1007/s00018-017-2740-5

Huberty, A. F., & Denno, R. F. (2004). Plant water stress and its consequences for herbivorous insects: A new synthesis. Ecology, 85(5), 1383–1398. https://doi.org/10.1890/03-0352

Hunter, M. D. (2016). The phytochemical landscape. Princeton, New Jersey: Princeton University Press.

Isah, T. (2019). Stress and defense responses in plant secondary metabolites production. Biological Research, 52(1), 39. https://doi.org/10.1186/s40659-019-0246-3

Islinger, M., Cardoso, M. J. R., & Schrader, M. (2010). Be different-The diversity of peroxisomes in the animal kingdom. Biochimica et Biophysica Acta - Molecular Cell Research, 1803(8), 881–897. https://doi.org/10.1016/j.bbamcr.2010.03.013

Jamieson, M. A., Trowbridge, A. M., Raffa, K. F., & Lindroth, R. L. (2012). Consequences of climate warming and altered precipitation patterns for plant-insect and multitrophic interactions. Plant Physiology, 160(4), 1719–1727. https://doi.org/10.1104/pp.112.206524

Jamieson, M. A., Burkle, L. A., Manson, J. S., Runyon, J. B., Trowbridge, A. M., & Zientek, J. (2017). Global change effects on plant–insect interactions: the role of phytochemistry. Current Opinion in Insect Science, 23, 70–80.https://doi.org/10.1016/j.cois.2017.07.009

Kahilainen, A., van Nouhuys, S., Schulz, T., & Saastamoinen, M. (2018). Metapopulation dynamics in a changing climate: Increasing spatial synchrony in weather conditions drives metapopulation synchrony of a butterfly inhabiting a fragmented landscape. Global Change Biology, 24(9), 4316–4329. https://doi.org/10.1111/gcb.14280

Kalinkat, G., Jochum, M., Brose, U., & Dell, A. I. (2015). Body size and the behavioral ecology of insects: Linking individuals to ecological communities. Current Opinion in Insect Science, 9, 24–30. https://doi.org/10.1016/j.cois.2015.04.017

Kim, H. K., Choi, Y. H., & Verpoorte, R. (2010). NMR-based metabolomic analysis of plants. Nature Protocols, 5(3), 536–549. https://doi.org/10.1038/nprot.2009.237

Kuglerová, M., Skuhrovec, J., & Münzbergová, Z. (2019). Relative importance of drought, soil quality, and plant species in determining the strength of plant–herbivore interactions. Ecological Entomology, een.12745. https://doi.org/10.1111/een.12745

Kuussaari, M., van Nouhuys, S., Hellemann, J. J., & Singer, M. C. (2004). Larval Biology of Checkerspots. In P. R. Ehrlich & I. Hanski (Eds.), On the Wings of Checkerspots: A Model System for Population Biology (pp. 138–160). New York: Oxford University Press.

Kvist, J., Wheat, C. W., Kallioniemi, E., Saastamoinen, M., Hanski, I., & Frilander, M. J. (2013). Temperature treatments during larval development reveal extensive heritable and plastic variation in gene expression and life history traits. Molecular Ecology, 22(3), 602–619. https://doi.org/10.1111/j.1365-294X.2012.05521.x

Langmead, B., Trapnell, C., Pop, M., & Salzberg, S. L. (2009). Ultrafast and memory-efficient alignment of short DNA sequences to the human genome. Genome Biology, 10(3), R25. https://doi.org/10.1186/gb-2009-10-3-r25

Larsson, S. (1989). Stressful times for the plant stress: Insect performance hypothesis. Oikos, 56(2), 277–283. https://doi.org/10.2307/3565348

Li, B., & Dewey, C. N. (2011). RSEM: accurate transcript quantification from RNA-Seq data with or without a reference genome. BMC Bioinformatics, 12(1), 323. https://doi.org/10.1186/1471-2105-12-323

Li, N., Liu, Q., Xiong, Y., & Yu, J. (2019). Headcase and Unkempt regulate tissue growth and cell cycle progression in response to nutrient restriction. Cell Reports, 26(3), 733–747. https://doi.org/10.1016/j.celrep.2018.12.086

Li, X., Garrity, A. G., & Xu, H. (2013, September). Regulation of membrane trafficking by signalling on endosomal and lysosomal membranes. Journal of Physiology. https://doi.org/10.1113/jphysiol.2013.258301

Mattson, W. J., & Haack, R. a. (1987). The role of drought in outbreaks of plant-eating insects - Drought’s physiological effects on plants can predict its influence on insect populations. BioScience, 37(2), 110–118.

McCarthy, D. J., Chen, Y., & Smyth, G. K. (2012). Differential expression analysis of multifactor RNA-Seq experiments with respect to biological variation. Nucleic Acids Research, 40(10), 4288–4297. https://doi.org/10.1093/nar/gks042

Medlar, A. J., Törönen, P., & Holm, L. (2018). AAI-profiler: fast proteome-wide exploratory analysis reveals taxonomic identity, misclassification and contamination. Nucleic Acids Research, 46(W1), W479–W485. https://doi.org/10.1093/nar/gky359

Nallu, S., Hill, J. A., Don, K., Sahagun, C., Zhang, W., Meslin, C., et al. (2018). The molecular genetic basis of herbivory between butterflies and their host plants. Nature Ecology and Evolution, 2(9), 1418–1427. https://doi.org/10.1038/s41559-018-0629-9

Nieminen, M., Siljander, M., & Hanski, I. (2004). Structure and Dynamics of Melitaea cinxia Metapopulations. In P. R. Ehrlich & I. Hanski (Eds.), On the Wings of Checkerspots: A Model System for Population Biology (pp. 63–91). New York: Oxford University Press.

Niitepõld, K., & Saastamoinen, M. (2017). A candidate gene in an ecological model species: Phosphoglucose isomerase (Pgi) in the Glanville Fritillary butterfly (Melitaea cinxia). Annales Zoologici Fennici, 54(1–4), 259–273. https://doi.org/10.5735/086.054.0122

Ojanen, S. P., Nieminen, M., Meyke, E., Pöyry, J., & Hanski, I. (2013). Long-term metapopulation study of the Glanville fritillary butterfly (Melitaea cinxia): Survey methods, data management, and long-term population trends. Ecology and Evolution, 3(11), 3713–3737. https://doi.org/10.1002/ece3.733

Oksanen, J., Blanchet, F. G., Friendly, M., Kindt, R., Legendre, P., McGlinn, D., et al. (2018). vegan: Community Ecology Package. Retrieved from https://cran.r-project.org/package=vegan

Oliver, M. J., Guo, L., Alexander, D. C., Ryals, J. A., Wone, B. W. M., & Cushman, J. C. (2011). A sister group contrast using untargeted global metabolomic analysis delineates the biochemical regulation underlying desiccation tolerance in Sporobolus stapfianus. The Plant Cell, 23(4), 1231–1248. https://doi.org/10.1105/tpc.110.082800

Pincebourde, S., van Baaren, J., Rasmann, S., Rasmont, P., Rodet, G., Martinet, B., & Calatayud, P.-A. (2017). Plant–insect interactions in a changing world. Advances in Botanical Research, 81, 289–332. https://doi.org/10.1016/BS.ABR.2016.09.009

Piper, R. C., & Lehner, P. J. (2011). Endosomal transport via ubiquitination. Trends in Cell Biology, 21(11), 647–655. https://doi.org/10.1016/j.tcb.2011.08.007

Post, E. (2013). Erosion of community diversity and stability by herbivore removal under warming. Proceedings of the Royal Society B: Biological Sciences, 280(1757), 20122722. https://doi.org/10.1098/rspb.2012.2722

Price, T. D., Qvarnström, A., & Irwin, D. E. (2003). The role of phenotypic plasticity in driving genetic evolution. Proceedings of the Royal Society of London. Series B: Biological Sciences, 270(1523), 1433–1440. https://doi.org/10.1098/rspb.2003.2372

Van der Putten, W. H., Macel, M., & Visser, M. E. (2010). Predicting species distribution and abundance responses to climate change: why it is essential to include biotic interactions across trophic levels. Philosophical Transactions of the Royal Society of London. Series B, Biological Sciences, 365(1549), 2025–2034. https://doi.org/10.1098/rstb.2010.0037

R Core Team. (2019). R: A language and environment for statistical computing. Vienna, Austria: R Foundation for Statistical Computing. Retrieved from http://www.r-project.org

Rizhsky, L., Liang, H., Shuman, J., Shulaev, V., Davletova, S., & Mittler, R. (2004). When defense pathways collide. The response of Arabidopsis to a combination of drought and heat stress. Plant Physiology, 134(April), 1683–1696. https://doi.org/10.1104/pp.103.033431.1

Robinson, M. D., & Oshlack, A. (2010). A scaling normalization method for differential expression analysis of RNA-seq data. Genome Biology, 11(3), R25. https://doi.org/10.1186/gb-2010-11-3-r25

Robinson, M. D., McCarthy, D. J., & Smyth, G. K. (2010). edgeR: a Bioconductor package for differential expression analysis of digital gene expression data. Bioinformatics, 26(1), 139–140. https://doi.org/10.1093/bioinformatics/btp616

Rosa, E., & Saastamoinen, M. (2017). Sex-dependent effects of larval food stress on adult performance under semi-natural conditions: only a matter of size? Oecologia, 184(3), 633–642. https://doi.org/10.1007/s00442-017-3903-7

Rosa, E., Minard, G., Lindholm, J., & Saastamoinen, M. (2019). Moderate plant water stress improves larval development, and impacts immunity and gut microbiota of a specialist herbivore. PLOS ONE, 14(2), e0204292. https://doi.org/10.1371/journal.pone.0204292

Rosenblatt, A. E., & Schmitz, O. J. (2016). Climate change, nutrition, and bottom-up and top-down food web processes. Trends in Ecology & Evolution, 31(12), 965–975. https://doi.org/10.1016/j.tree.2016.09.009

Saastamoinen, M., Ikonen, S., Wong, S. C., Lehtonen, R., & Hanski, I. (2013). Plastic larval development in a butterfly has complex environmental and genetic causes and consequences for population dynamics. Journal of Animal Ecology, 82(3), 529–539. https://doi.org/10.1111/1365-2656.12034

Salgado, A. L., & Saastamoinen, M. (2019). Developmental stage-dependent response and preference for host plant quality in an insect herbivore. Animal Behaviour, 150, 27–38. https://doi.org/10.1016/j.anbehav.2019.01.018

Schlicker, A., Domingues, F. S., Rahnenführer, J., & Lengauer, T. (2006). A new measure for functional similarity of gene products based on Gene Ontology. BMC Bioinformatics, 7, 302. https://doi.org/10.1186/1471-2105-7-302

Schulz, M. H., Zerbino, D. R., Vingron, M., & Birney, E. (2012). Oases: robust *de novo* RNA-seq assembly across the dynamic range of expression levels. Bioinformatics, 28(8), 1086–1092. https://doi.org/10.1093/bioinformatics/bts094

Scott, R. C., Schuldiner, O., & Neufeld, T. P. (2004). Role and regulation of starvation-induced autophagy in the Drosophila fat body. Developmental Cell, 7(2), 167–178. https://doi.org/10.1016/j.devcel.2004.07.009

Seppey, M., Ioannidis, P., Emerson, B. C., Pitteloud, C., Robinson-Rechavi, M., Roux, J., et al. (2019). Genomic signatures accompanying the dietary shift to phytophagy in polyphagan beetles. Genome Biology, 20(1), 98. https://doi.org/10.1186/s13059-019-1704-5

Siepielski, A. M., Hasik, A. Z., & Ousterhout, B. H. (2018). An ecological and evolutionary perspective on species coexistence under global change. Current Opinion in Insect Science, 29, 71–77. https://doi.org/10.1016/j.cois.2018.06.007

Sonoda, S., Fukumoto, K., Izumi, Y., Ashfaq, M., Yoshida, H., & Tsumuki, H. (2007). Expression profile of arylphorin gene during diapause and cold acclimation in the rice stem borer, Chilo suppressalis Walker (Lepidoptera: Crambidae). Applied Entomology and Zoology, 42(1), 35–40. https://doi.org/10.1303/aez.2007.35

Stan Development Team. (2018). RStan: the R interface to Stan. Retrieved from http://mc-stan.org/

Stange, M., Barrett, R. D. H., & Hendry, A. P. (2020). The importance of genomic variation for biodiversity, ecosystems and people. Nature Reviews Genetics, 22(2), 89–105. https://doi.org/10.1038/s41576-020-00288-7

Tack, A. J. M., Mononen, T., & Hanski, I. (2015). Increasing frequency of low summer precipitation synchronizes dynamics and compromises metapopulation stability in the Glanville fritillary butterfly. Proceedings of the Royal Society B: Biological Sciences, 282, 20150173. https://doi.org/10.1098/rspb.2015.0173

The Gene Ontology Consortium, Ashburner, M., Ball, C. A., Blake, J. A., Botstein, D., Butler, H., et al. (2000). Gene Ontology: tool for the unification of biology. Nature Genetics, 25(1), 25. https://doi.org/10.1038/75556

Törönen, P., Medlar, A., & Holm, L. (2018). PANNZER2: a rapid functional annotation web server. Nucleic Acids Research, 46(W1), W84–W88. https://doi.org/10.1093/nar/gky350

Tylianakis, J. M., Didham, R. K., Bascompte, J., & Wardle, D. A. (2008). Global change and species interactions in terrestrial ecosystems. Ecology Letters, 11(12), 1351–1363. https://doi.org/10.1111/j.1461-0248.2008.01250.x

Verspagen, N., Ikonen, S., Saastamoinen, M., & van Bergen, E. (2020). Multidimensional plasticity in the Glanville fritillary butterfly: larval performance is temperature, host and family specific. Proceedings of the Royal Society B: Biological Sciences, 287(1941), 20202577. https://doi.org/10.1098/rspb.2020.2577

Via, S., & Lande, R. (1985). Genotype-environment interaction and the evolution of phenotypic plasticity. Evolution, 39(3), 505–522.

Vogel, H., Musser, R. O., & Celorio-Mancera, M. de la P. (2014). Transcriptome Responses in Herbivorous Insects Towards Host Plant and Toxin Feeding. In C. Voelckel & G. Jander (Eds.), Annual Plant Reviews: Insect-Plant Interactions, Volume 47 (pp. 197– 233). Chichester, UK: John Wiley & Sons, Ltd. https://doi.org/10.1002/9781118829783.ch6

Voigt, W., Perner, J., Davis, A. J., Eggers, T., Schumacher, J., Bährmann, R., et al. (2003). Trophic levels are differentially sensitive to climate. Ecology, 84(9), 2444–2453. https://doi.org/10.1890/02-0266

Wahlberg, N. (2000). Comparative descriptions of immature stages of five Finnish melitaeine butterfly species (Lepidoptera: Nyphalidae). Entomologica Fennica, 11, 167–173.

Walter, J., Hein, R., Auge, H., Beierkuhnlein, C., Löffler, S., Reifenrath, K., et al. (2012). How do extreme drought and plant community composition affect host plant metabolites and herbivore performance? Arthropod-Plant Interactions, 6(1), 15–25. https://doi.org/10.1007/s11829-011-9157-0

Waring, G. L., & Cobb, N. S. (1992). The impact of plant stress on herbivore population dynamics. In E. Bernays (Ed.), Insect-Plant Interactions (1st ed., pp. 167–226). Boca Raton: CRC Press. https://doi.org/10.1201/9781351271004

Weisser, W. W., & Siemann, E. (Eds.). (2008). Insects and Ecosystem Function (Vol. 173). Berlin, Heidelberg: Springer Berlin Heidelberg. https://doi.org/10.1007/978-3-540-74004-9

Wennersten, L., & Forsman, A. (2012). Population-level consequences of polymorphism, plasticity and randomized phenotype switching: A review of predictions. Biological Reviews, 87(3), 756–767. https://doi.org/10.1111/j.1469-185X.2012.00231.x

White, T. C. R. (1974). A hypothesis to explain outbreaks of looper caterpillars, with special reference to populations of Selidosema suavis in a plantation of Pinus radiata in New Zealand. Oecologia, 16(4), 279–301. Retrieved from https://www.jstor.org/stable/4215010?seq=1#metadata_info_tab_contents

White, T. C. R. (2009). Plant vigour versus plant stress: a false dichotomy. Oikos, 118(6), 807–808. https://doi.org/10.1111/j.1600-0706.2009.17495.x

Wirta, H. K., Vesterinen, E. J., Hambäck, P. A., Weingartner, E., Rasmussen, C., Reneerkens, J., et al. (2015). Exposing the structure of an Arctic food web. Ecology and Evolution, 5(17), 3842–3856. https://doi.org/10.1002/ece3.1647

Wu, T. D., & Watanabe, C. K. (2005). GMAP: a genomic mapping and alignment program for mRNA and EST sequences. Bioinformatics, 21(9), 1859–1875. https://doi.org/10.1093/bioinformatics/bti310

Wullschleger, S., Loewith, R., & Hall, M. N. (2006, February 10). TOR signaling in growth and metabolism. Cell. https://doi.org/10.1016/j.cell.2006.01.016

Yang, L. H., Cenzer, M. L., Morgan, L. J., & Hall, G. W. (2020). Species-specific, age-varying plant traits affect herbivore growth and survival. Ecology, 101(7), e03029. https://doi.org/10.1002/ecy.3029

Zhang, H., Stallock, J. P., Ng, J. C., Reinhard, C., & Neufeld, T. P. (2000). Regulation of cellular growth by the Drosophila target of rapamycin dTOR. Genes and Development, 14(21), 2712–2724. https://doi.org/10.1101/gad.835000

