## Supplementary figures & tables for "Alternative developmental and transcriptomic responses to host plant water limitation in a butterfly metapopulation"

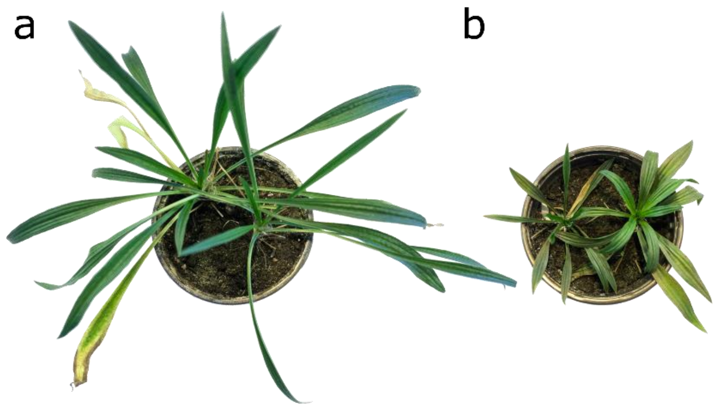


**Figure S1.** Typical examples of (a) control and (b) water limited *P. lanceolata*.

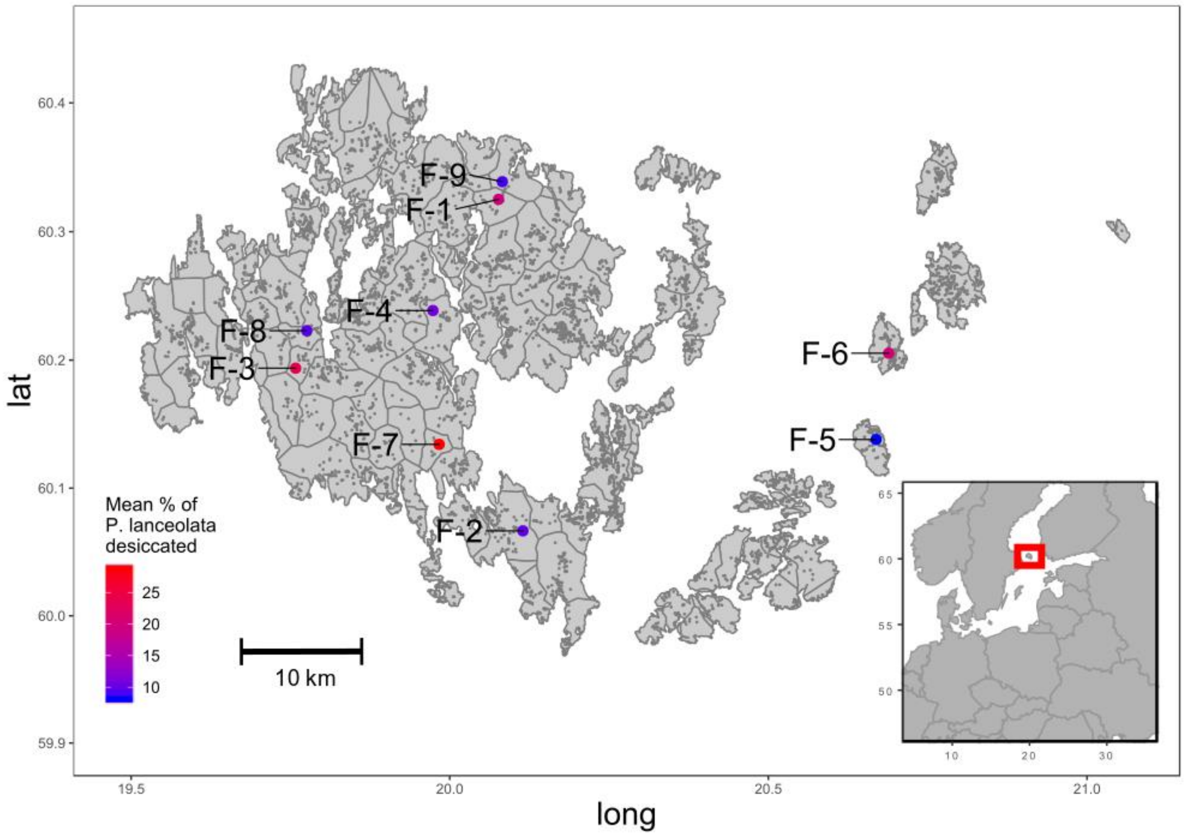


**Figure S2.** Sampling localities of the parents used to breed larval full-sib families. Sampling localities marked with red dots, grey dots and polygons represent habitat patches and their division into semi-independent networks (SINs), respectively.

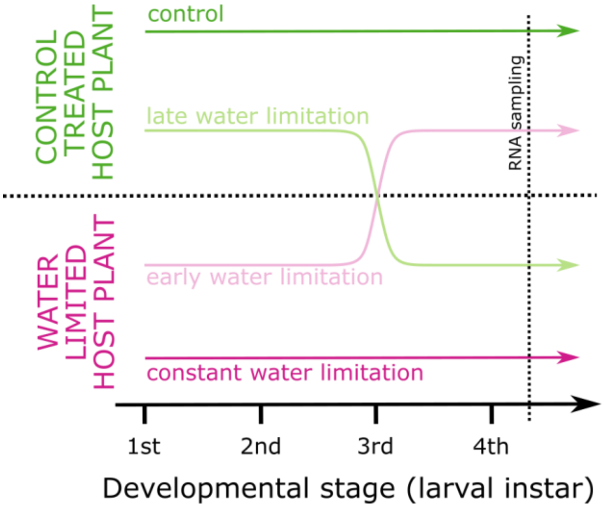


**Figure S3.** Experimental treatments. The coloured lines indicate feeding on control or drought stressed host plant leaf tissue during different pre-diapause larval instars in the different experimental treatments.

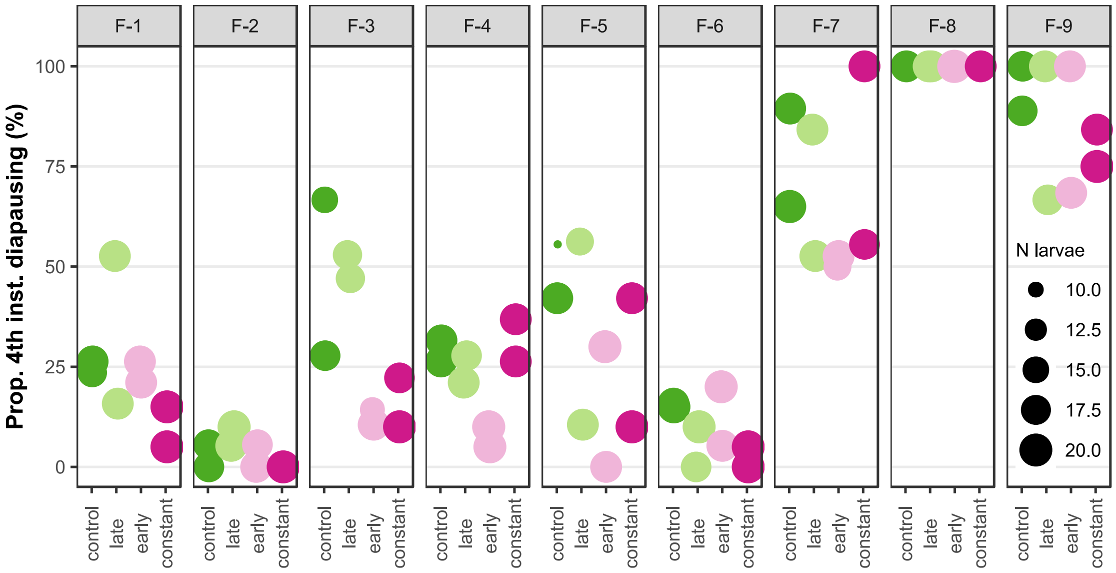


**Figure S4.** Proportion of larvae entering diapause in the 4$^{th}$ larval instar per family per treatment. Although most families exhibited both diapause strategies, the division between primarily 5th instar and 4th instar diapausing families is rather clear-cut as 4th instar diapause occurred typically in less than 30% of the larvae in the primarily 5th instar diapausing families and in more than 70% in the primarily 4th instar diapausing families.

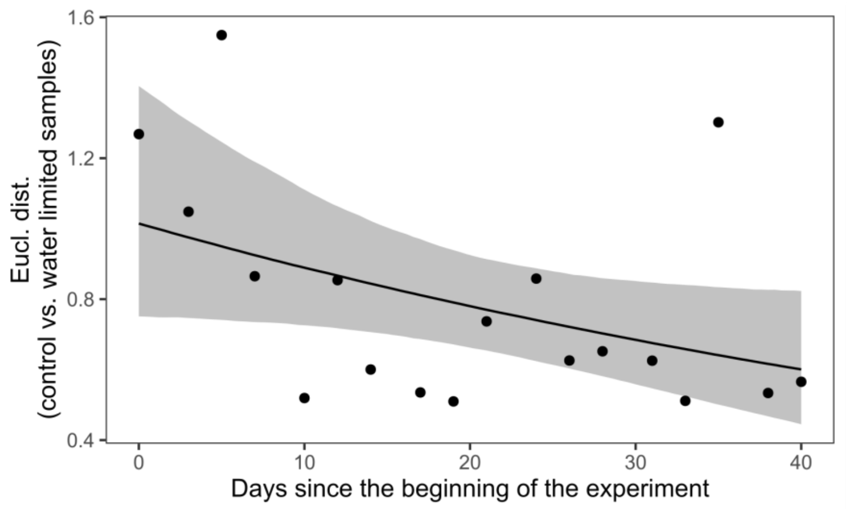


**Figure S5.** Convergence of the metabolomes of the control and drought treated plants with time. Euclidean distance between control and water limited plants. The points represent the observed distances, and the solid black line with the grey shading represent the trend and its 95% credible interval as modelled in a Gamma distribution GLM (see methods).

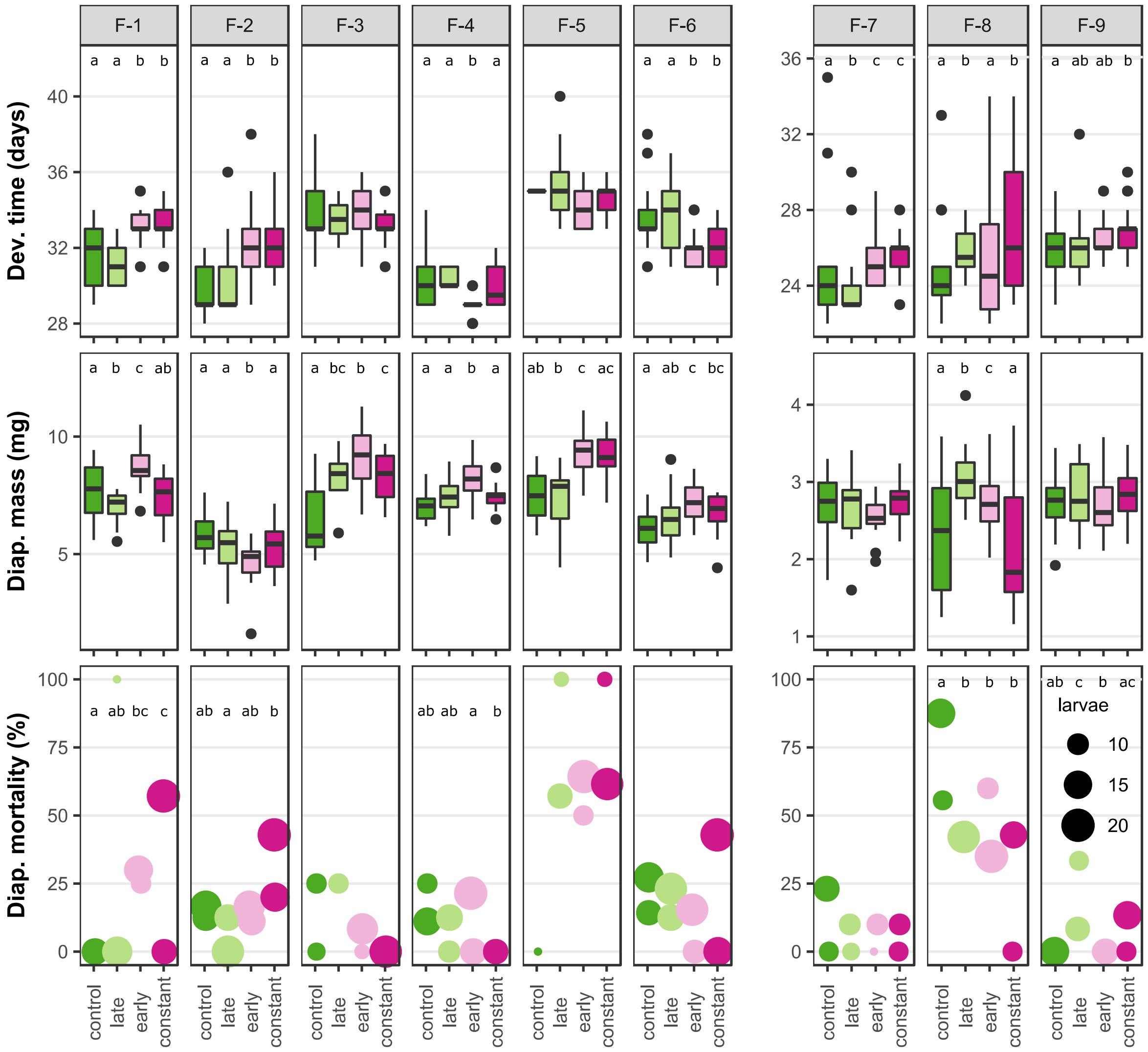


**Figure S6.** Development time, diapause body mass and overwintering mortality per family per treatment. The lower case letters on top of each strip illustrates pairwise differences between treatment levels. Note that control and late treatment levels on one hand and early and constant on the other often group together.

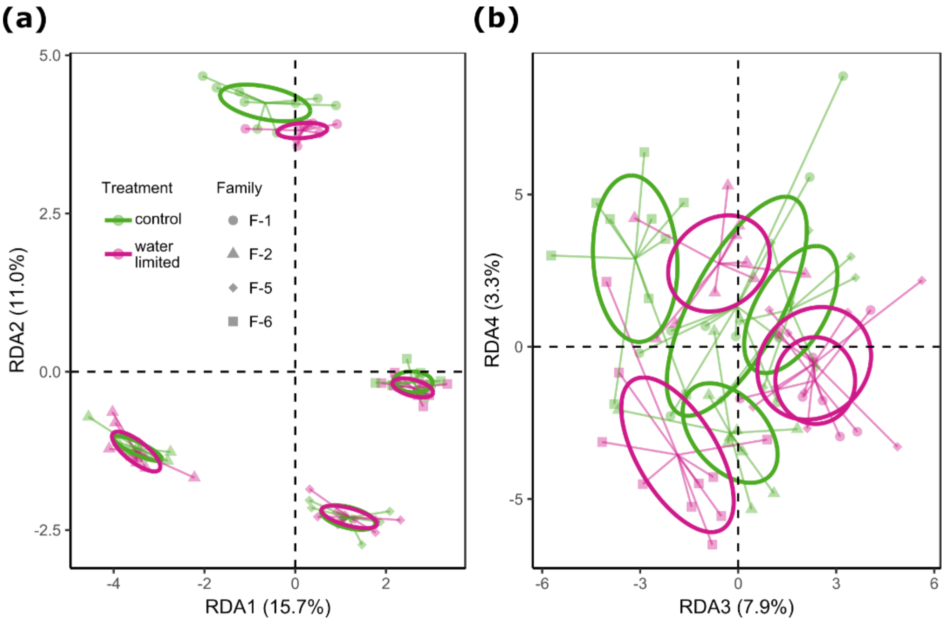


**Figure S7.** RDA examining the associations between family, treatment (control treatment combined with late drought and early drought with constant drought) and their interaction with the transcriptome of *M. cinxia* larvae. The individual scores (points), group centroids (the centre of line spiders) and the 95% confidence interval of the centroids (ellipses) along four statistically significant constrained axes (a: RDA1 and RDA2; b: RDA3 and RDA4.

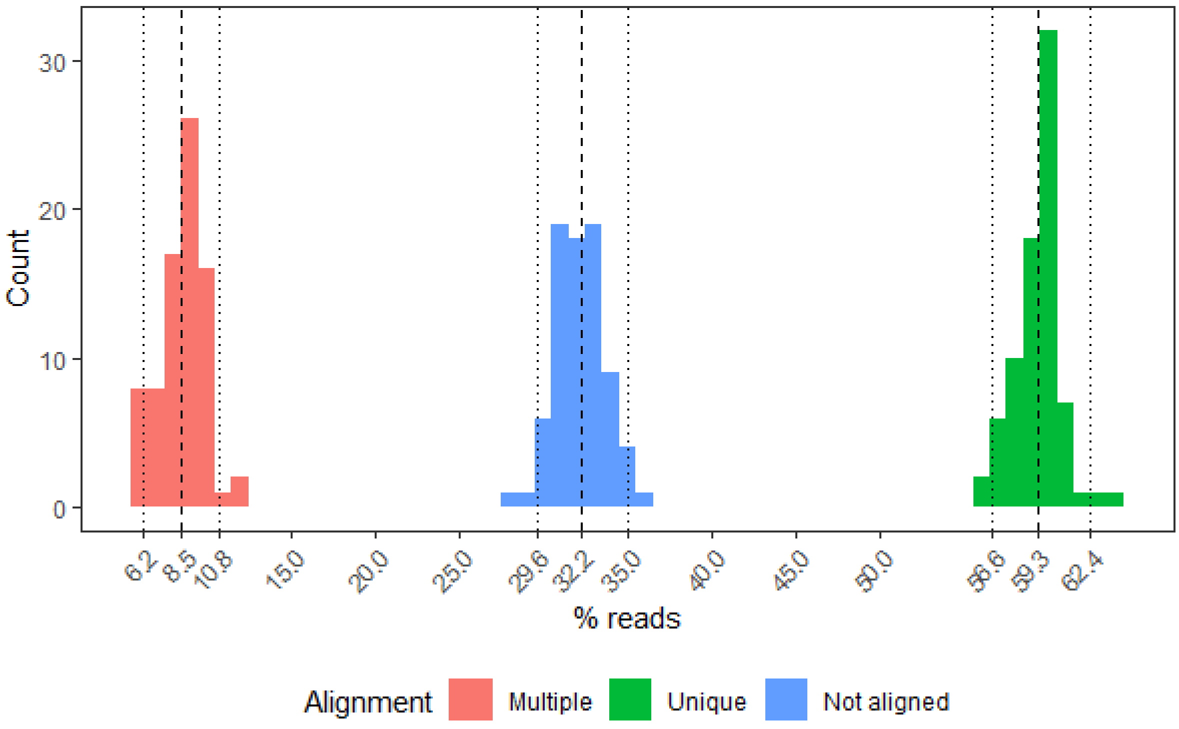


**Figure S8.** Mapping of reads on the de novo transcriptome. Dashed lines represent mean percentage and dotted lines the 95% interval for each category.

**Table S1.** Characteristics of the sampling localities of the parent individuals used to create full-sib family groups, and description of the percentage of offspring entering diapause in the 4$^{th}$ larval instar.

| Family ID | Lat. | Long. | Mean % of *P. lanceolata* desiccated | % 4th instar diapause |
|---|---|---|---|---|
| F-1 | 60.33 | 20.08 | 19.41 | 22.4 |
| F-2 | 60.07 | 20.11 | 9.91 | 3.3 |
| F-3 | 60.19 | 19.76 | 24.00 | 30.2 |
| F-4 | 60.24 | 19.97 | 10.90 | 22.9 |
| F-5 | 60.14 | 20.67 | 8.06 | 28.4 |
| F-6 | 60.21 | 20.69 | 20.47 | 9.0 |
| F-7 | 60.13 | 19.98 | 28.85 | 69.1 |
| F-8 | 60.22 | 19.78 | 9.56 | 100.0 |
| F-9 | 60.34 | 20.08 | 9.06 | 83.8 |

**Table S2.** The numbers of larvae sampled, Sequenom panel genotyped for sexing and transcriptome sequenced per family per egg clutch per treatment.

| Family ID | Treatment | $N_{collected}$ | $N_{sequenom}$ | $N_{RNA\text{-}seq}$ |
|---|---|---|---|---|
| F-1 | Control | 10+5 | 9+5 | 3+2 |
| | Late water lim. | 10+5 | 8+5 | 2+2 |
| | Early water lim. | 10+5 | 10+5 | 3+2 |
| | Constant water lim. | 10+5 | 10+5 | 4+1 |
| F-2 | Control | 10+5 | 10+5 | 3+2 |
| | Late water lim. | 10+5 | 10+5 | 3+2 |
| | Early water lim. | 10+5 | 10+5 | 3+2 |
| | Constant water lim. | 10+5 | 10+5 | 3+2 |
| F-3 | Control | 10+2 | | |
| | Late water lim. | 10+5 | | |
| | Early water lim. | 10+5 | | |
| | Constant water lim. | 10+5 | | |
| F-4 | Control | 10+5 | | |
| | Late water lim. | 10+5 | | |
| | Early water lim. | 10+5 | | |
| | Constant water lim. | 10+5 | | |
| F-5 | Control | 5+10 | 0+10 | 0+4 |
| | Late water lim. | 10+10 | 5+10 | 3+2 |
| | Early water lim. | 10+5 | 10+5 | 4+1 |
| | Constant water lim. | 10+5 | 9+5 | 3+2 |
| F-6 | Control | 10+5 | 10+5 | 3+2 |
| | Late water lim. | 10+5 | 10+5 | 3+2 |
| | Early water lim. | 10+5 | 10+5 | 3+2 |
| | Constant water lim. | 10+5 | 10+5 | 5+0 |
| F-7 | Control | 10 | | |
| | Late water lim. | 10 | | |
| | Early water lim. | 10 | | |
| | Constant water lim. | 10 | | |
| F-8 | Control | 10 | | |
| | Late water lim. | 10 | | |
| | Early water lim. | 10 | | |
| | Constant water lim. | 10 | | |
| F-9 | Control | 10 | | |
| | Late water lim. | 10 | | |
| | Early water lim. | 10 | | |
| | Constant water lim. | 10 | | |

**Table S3.** Model coefficient estimates (Est. coef.), their standard errors (SE) and 95% credible intervals (95% Cr. I.) for the effect of time on the Euclidean distance between the metabolomes of control and water limited *P. lanceolata*.

| **Covariate** | **Est. coef.** | **SE** | **95% Cr.I.** | |
|---|---|---|---|---|
| | | | Lower | Upper |
| Intercept | 0.017 | 0.158 | -0.286 | 0.340 |
| Time | -0.013 | 0.007 | -0.026 | 0.000 |

**Table S4.** Model coefficient estimates (Est.), their standard errors (SE) and 95% credible intervals (95% Cr. I.) for the effects of family identity, water limitation treatment and their interaction on the different developmental responses of *M. cinxia.* Water limitation treatment refers to feeding on water limited *P. lanceolata* during the first two instars of their pre-diapause development (early & constant water limitation combined for water limitation treatment and control & late water limitation combined for control treatment; Fig. S3).

| | **Development time** | | | | **Diapause mass** | | | | **OW mortality** | | | |
|---|---|---|---|---|---|---|---|---|---|---|---|---|
| | **Est.** | **SE** | **95% Cr.I.** | | **Est.** | **SE** | **95% Cr.I.** | | **Est.** | **SE** | **95% Cr.I.** | |
| **Coefficient** | | | L | U | | | L | U | | | L | U |
| *5th instar diapausing larvae* | | | | | | | | | | | | |
| Intercept [illegible] | **1.92** | 0.15 | 1.63 | 2.22 | **7.30** | 0.39 | 6.53 | 8.05 | **-2.90** | 0.91 | -4.80 | -1.23 |
| F-2 | **-0.21** | 0.11 | -0.43 | -0.01 | **-1.74** | 0.53 | -2.78 | -0.69 | 0.48 | 1.18 | -1.88 | 2.84 |
| F-3 | **0.30** | 0.11 | 0.08 | 0.53 | -0.18 | 0.60 | -1.34 | 0.99 | 1.00 | 1.36 | -1.77 | 3.57 |
| F-4 | -0.15 | 0.11 | -0.36 | 0.06 | -0.06 | 0.55 | -1.13 | 1.04 | 0.51 | 1.24 | -1.99 | 2.98 |
| F-5 | **0.48** | 0.12 | 0.25 | 0.74 | -0.26 | 0.62 | -1.47 | 0.89 | **3.33** | 1.24 | 0.92 | 5.85 |
| F-6 | **0.29** | 0.11 | 0.09 | 0.51 | -0.90 | 0.54 | -1.97 | 0.17 | 1.24 | 1.15 | -1.07 | 3.51 |
| Water limitation | **0.26** | 0.05 | 0.16 | 0.37 | **0.56** | 0.27 | 0.03 | 1.09 | **2.10** | 0.81 | 0.62 | 3.80 |
| F-2 x Water limitation | 0.06 | 0.06 | -0.05 | 0.19 | **-1.15** | 0.35 | -1.83 | -0.46 | -0.87 | 1.02 | -2.90 | 1.08 |
| F-3 x Water limitation | **-0.31** | 0.08 | -0.48 | -0.15 | **1.01** | 0.45 | 0.13 | 1.89 | **-3.99** | 1.57 | -7.26 | -1.07 |
| F-4 x Water limitation | **-0.45** | 0.09 | -0.64 | -0.29 | 0.17 | 0.38 | -0.58 | 0.92 | **-2.29** | 1.19 | -4.68 | -0.03 |
| F-5 x Water limitation | **-0.34** | 0.09 | -0.53 | -0.19 | **1.38** | 0.46 | 0.48 | 2.29 | -1.85 | 1.10 | -4.07 | 0.22 |
| F-6 x Water limitation | **-0.44** | 0.08 | -0.60 | -0.29 | 0.10 | 0.36 | -0.60 | 0.79 | **-2.20** | 0.99 | -4.20 | -0.36 |
| $\sigma_{(family:clutch)}$ | 0.08 | 0.04 | 0.03 | 0.18 | 0.39 | 0.24 | 0.07 | 0.95 | 0.71 | 0.45 | 0.06 | 1.82 |
| $\sigma_{res}$ | 0.17 | 0.02 | 0.13 | 0.22 | 1.03 | 0.04 | 0.96 | 1.11 | | | | |
| Dist. shift | 24.30 | 0.93 | 22.17 | 25.77 | | | | | | | | |
| *4th instar diapausing larvae* | | | | | | | | | | | | |
| Intercept [illegible] | **1.44** | 0.33 | 0.80 | 2.04 | **2.67** | 0.47 | 2.02 | 3.33 | **-2.71** | 1.37 | -5.11 | -0.87 |
| F-8 | 0.22 | 0.43 | -0.59 | 1.03 | 0.11 | 0.64 | -0.79 | 1.01 | **2.80** | 1.69 | 0.18 | 5.55 |
| F-9 | 0.37 | 0.43 | -0.43 | 1.18 | 0.13 | 0.64 | -0.80 | 1.03 | 0.21 | 1.74 | -2.36 | 3.13 |
| Water limitation | **0.29** | 0.07 | 0.16 | 0.44 | -0.01 | 0.11 | -0.19 | 0.16 | -0.76 | 0.94 | -2.36 | 0.73 |
| F-8 x Water limitation | **-0.26** | 0.09 | -0.45 | -0.10 | -0.20 | 0.14 | -0.43 | 0.03 | -0.18 | 1.01 | -1.78 | 1.55 |
| F-9 x Water limitation | **-0.16** | 0.08 | -0.33 | 0.00 | -0.01 | 0.14 | -0.25 | 0.23 | -0.13 | 1.31 | -2.27 | 2.02 |
| $\sigma_{(family:clutch)}$ | 0.32 | 0.30 | 0.09 | 1.10 | 0.53 | 0.44 | 0.19 | 1.29 | 1.45 | 1.38 | 0.12 | 4.08 |
| $\sigma_{res}$ | 0.31 | 0.03 | 0.24 | 0.37 | 0.46 | 0.02 | 0.42 | 0.49 | | | | |
| Dist. shift | 19.66 | 0.55 | 18.39 | 20.54 | | | | | | | | |

**Table S5.** Model coefficient estimates (Est.), their standard errors (SE) and 95% credible intervals (95% Cr. I.) for the effects of family, treatment and their interaction on developmental responses of *M. cinxia*, while maintaining all original treatment levels (Figure S3). Estimates with a credible interval differing from zero are bolded.

| | **Development time** | | | | **Diapause mass** | | | | **OW mortality** | | | |
|---|---|---|---|---|---|---|---|---|---|---|---|---|
| **Coefficient** | **Est.** | **SE** | **95% Cr.I.** | | **Est.** | **SE** | **95% Cr.I.** | | **Est.** | **SE** | **95% Cr.I.** | |
| | | | L | U | | | L | U | | | L | U |
| *5th instar diapausing larvae* | | | | | | | | | | | | |
| Intercept [illegible] | **1.75** | 0.14 | 1.47 | 2.03 | **7.66** | 0.48 | 6.71 | 8.61 | **-2.88** | 1.02 | -5.02 | -1.03 |
| F-2 | **-0.35** | 0.14 | -0.64 | -0.08 | **-1.89** | 0.67 | -3.22 | -0.57 | 0.84 | 1.33 | -1.77 | 3.53 |
| F-3 | **0.36** | 0.15 | 0.06 | 0.67 | **-1.16** | 0.71 | -2.54 | 0.25 | 0.12 | 1.65 | -3.30 | 3.25 |
| F-4 | -0.24 | 0.14 | -0.52 | 0.03 | -0.59 | 0.69 | -1.95 | 0.77 | 0.52 | 1.45 | -2.35 | 3.34 |
| F-5 | **0.51** | 0.25 | 0.03 | 1.02 | -0.18 | 0.92 | -2.00 | 1.60 | 0.64 | 2.19 | -3.79 | 4.77 |
| F-6 | **0.31** | 0.14 | 0.04 | 0.59 | **-1.53** | 0.67 | -2.84 | -0.21 | 1.18 | 1.31 | -1.38 | 3.77 |
| Late | -0.09 | 0.09 | -0.26 | 0.08 | **-0.83** | 0.40 | -1.61 | -0.05 | -0.05 | 1.22 | -2.56 | 2.26 |
| Early | **0.27** | 0.08 | 0.11 | 0.44 | **0.96** | 0.38 | 0.20 | 1.69 | 1.62 | 1.01 | -0.24 | 3.66 |
| Constant | **0.27** | 0.08 | 0.12 | 0.43 | -0.35 | 0.35 | -1.03 | 0.34 | **2.13** | 0.94 | 0.44 | 4.12 |
| F-2 x Late | 0.17 | 0.11 | -0.04 | 0.39 | 0.43 | 0.50 | -0.58 | 1.44 | -1.44 | 1.76 | -5.06 | 1.81 |
| F-3 x Late | -0.05 | 0.16 | -0.37 | 0.27 | **2.64** | 0.76 | 1.16 | 4.11 | 1.37 | 2.15 | -2.75 | 5.50 |
| F-4 x Late | 0.12 | 0.12 | -0.11 | 0.36 | **1.17** | 0.55 | 0.07 | 2.24 | -0.69 | 1.83 | -4.39 | 2.80 |
| F-5 x Late | 0.11 | 0.24 | -0.36 | 0.57 | 0.19 | 0.88 | -1.51 | 1.91 | 3.10 | 2.27 | -1.20 | 7.77 |
| F-6 x Late | 0.08 | 0.11 | -0.13 | 0.29 | **1.34** | 0.51 | 0.34 | 2.33 | -0.09 | 1.45 | -2.88 | 2.78 |
| F-2 x Early | 0.15 | 0.11 | -0.06 | 0.36 | **-2.09** | 0.48 | -3.02 | -1.15 | -1.57 | 1.32 | -4.18 | 1.00 |
| F-3 x Early | **-0.34** | 0.13 | -0.61 | -0.09 | **1.86** | 0.61 | 0.65 | 3.04 | -1.77 | 1.92 | -5.60 | 1.95 |
| F-4 x Early | **-0.60** | 0.13 | -0.87 | -0.36 | 0.23 | 0.51 | -0.78 | 1.23 | -1.40 | 1.44 | -4.14 | 1.38 |
| F-5 x Early | **-0.39** | 0.23 | -0.85 | 0.06 | 0.59 | 0.83 | -1.01 | 2.21 | 1.21 | 2.15 | -2.75 | 5.54 |
| F-6 x Early | **-0.51** | 0.11 | -0.74 | -0.30 | 0.26 | 0.50 | -0.71 | 1.23 | -2.59 | 1.38 | -5.39 | 0.06 |
| F-2 x Constant | **0.19** | 0.10 | 0.00 | 0.40 | -0.13 | 0.45 | -1.02 | 0.76 | -0.87 | 1.20 | -3.18 | 1.41 |
| F-3 x Constant | **-0.40** | 0.13 | -0.66 | -0.17 | **2.21** | 0.56 | 1.12 | 3.31 | **-5.76** | 2.96 | -12.35 | -0.93 |
| F-4 x Constant | **-0.33** | 0.12 | -0.57 | -0.11 | 0.72 | 0.52 | -0.30 | 1.73 | **-5.82** | 2.85 | -12.31 | -1.22 |
| F-5 x Constant | -0.31 | 0.23 | -0.77 | 0.14 | **1.62** | 0.81 | 0.03 | 3.21 | 0.99 | 2.13 | -2.96 | 5.26 |
| F-6 x Constant | **-0.47** | 0.11 | -0.69 | -0.27 | **1.06** | 0.46 | 0.13 | 1.96 | -1.80 | 1.17 | -4.17 | 0.42 |
| $\sigma_{(family:clutch)}$ | 0.10 | 0.05 | 0.04 | 0.23 | **0.50** | 0.26 | 0.17 | 1.14 | 0.80 | 0.49 | 0.09 | 1.95 |
| $\sigma_{res}$ | 0.21 | 0.02 | 0.17 | 0.25 | **0.97** | 0.04 | 0.90 | 1.05 | | | | |
| Dist. shift | 25.57 | 0.57 | 24.28 | 26.50 | | | | | | | | |
| *4th instar diapausing larvae* | | | | | | | | | | | | |
| Intercept [illegible] | **1.34** | 0.36 | 0.63 | 2.00 | **2.67** | 0.48 | 1.70 | 3.59 | **-2.66** | 1.52 | -6.19 | -0.03 |
| F-8 | 0.02 | 0.49 | -0.88 | 0.94 | -0.28 | 0.68 | -1.61 | 1.09 | 3.50 | 1.85 | -0.27 | 7.47 |
| F-9 | 0.33 | 0.49 | -0.59 | 1.28 | 0.11 | 0.67 | -1.17 | 1.50 | -2.51 | 2.39 | -7.19 | 2.38 |
| Late | **-0.17** | 0.09 | -0.35 | 0.00 | -0.02 | 0.13 | -0.27 | 0.24 | -0.90 | 1.19 | -3.50 | 1.25 |
| Early | **0.25** | 0.10 | 0.06 | 0.46 | -0.13 | 0.13 | -0.40 | 0.13 | -1.25 | 1.32 | -4.18 | 1.05 |
| Constant | **0.27** | 0.09 | 0.10 | 0.45 | 0.08 | 0.13 | -0.17 | 0.33 | -1.03 | 1.22 | -3.66 | 1.20 |
| F-8 x Late | **0.48** | 0.12 | 0.24 | 0.73 | **0.78** | 0.17 | 0.44 | 1.12 | -0.59 | 1.33 | -3.04 | 2.23 |
| F-9 x Late | 0.21 | 0.12 | -0.03 | 0.46 | 0.08 | 0.18 | -0.26 | 0.43 | **4.61** | 2.07 | 0.88 | 8.99 |
| F-8 x Early | **-0.27** | 0.13 | -0.54 | -0.02 | **0.59** | 0.17 | 0.26 | 0.93 | -0.23 | 1.43 | -2.76 | 2.83 |
| F-9 x Early | -0.09 | 0.13 | -0.36 | 0.16 | 0.08 | 0.18 | -0.27 | 0.43 | -2.34 | 3.78 | -10.51 | 4.14 |
| F-8 x Constant | 0.15 | 0.12 | -0.08 | 0.38 | -0.22 | 0.17 | -0.56 | 0.11 | -1.24 | 1.37 | -3.79 | 1.57 |
| F-9 x Constant | -0.09 | 0.12 | -0.33 | 0.14 | -0.02 | 0.17 | -0.35 | 0.32 | 3.60 | 2.13 | -0.35 | 8.05 |
| $\sigma_{(family:clutch)}$ | 0.37 | 0.33 | 0.11 | 1.24 | 0.55 | 0.44 | 0.18 | 1.70 | 1.57 | 1.46 | 0.07 | 5.41 |
| $\sigma_{res}$ | 0.33 | 0.03 | 0.27 | 0.39 | 0.40 | 0.02 | 0.37 | 0.44 | | | | |
| Dist. shift | 20.31 | 0.37 | 19.49 | 20.91 | | | | | | | | |

**Table S6.** Results of RDA on the transcriptomic responses of *M. cinxia* larvae across families and all treatment levels (no levels combined).

| | Constraint/RDA | df | Variance | Pseudo-F | P |
|---|---|---|---|---|---|
| Constraint | Family | 3 | 847.01 | 14.598 | <0.001 |
| | Treatment | 3 | 85.57 | 1.475 | 0.019 |
| | Family*Treatment | 9 | 272.83 | 60.19 | <0.001 |
| | Residual | 61 | 1179.82 | | |
| RDA | RDA1 | 1 | 331.61 | 18.680 | <0.001 |
| | RDA2 | 1 | 232.10 | 13.075 | <0.001 |
| | RDA3 | 1 | 166.13 | 9.359 | <0.001 |
| | RDA4 | 1 | 88.90 | 5.001 | <0.001 |
| | Residual | 60 | 1065.13 | | |

**Table S7.** GO enrichment analysis results on biological processes enriched in transcripts associated with RDA1 and RDA4.

| RDA | GO ID | Term | $N_{annotated}$ | $N_{associated}$ | $P_{Fisher.elim}$ |
|---|---|---|---|---|---|
| RDA1 | GO:0035721 | intraciliary retrograde transport | 10 | 4 | <0.001 |
| | GO:0006030 | chitin metabolic process | 55 | 7 | <0.001 |
| | GO:0006313 | transposition, DNA-mediated | 41 | 6 | <0.001 |
| | GO:0042737 | drug catabolic process | 21 | 4 | 0.001 |
| | GO:0007018 | microtubule-based movement | 83 | 10 | 0.004 |
| | GO:0000255 | allantoin metabolic process | 5 | 2 | 0.005 |
| | GO:0007304 | chorion-containing eggshell formation | 6 | 2 | 0.007 |
| | GO:0055076 | transition metal ion homeostasis | 7 | 2 | 0.010 |
| RDA4 | GO:0048015 | phosphatidylinositol-mediated signaling | 13 | 5 | <0.001 |
| | GO:0070588 | calcium ion transmembrane transport | 27 | 7 | <0.001 |
| | GO:0110010† | basolateral protein secretion | 5 | 3 | 0.002 |
| | GO:0110011† | regulation of basement membrane organiza... | 5 | 3 | 0.002 |
| | GO:0061864† | basement membrane constituent secretion | 5 | 3 | 0.002 |
| | GO:0007264 | small GTPase mediated signal transductio... | 71 | 11 | 0.002 |
| | GO:0046854 | phosphatidylinositol phosphorylation | 24 | 6 | 0.002 |
| | GO:0036092 | phosphatidylinositol-3-phosphate biosynt... | 6 | 3 | 0.003 |
| | GO:0032507 | maintenance of protein location in cell | 12 | 4 | 0.003 |
| | GO:0034765 | regulation of ion transmembrane transpor... | 31 | 6 | 0.006 |
| | GO:0051453 | regulation of intracellular pH | 8 | 3 | 0.008 |

† Semantic similarity could not be calculated for these terms, but as the terms had an identical set of transcripts associated with them, they are considered synonymous in the present paper.

**Table S8.** The GO biological process annotations and loadings on RDA1 of the transcripts that contribute to the enrichment of the biological processes along RDA1 in table S7.

| Transcript ID | GO BP annotation | GO BP enrichment contribution | Load. |
|---|---|---|---|
| evgtrinLocDN37403c1g2t3 | intraciliary retrograde transport | intraciliary retrograde transport; microtubule-based movement | 0.250 |
| evgtrinLocDN24084c0g1t3 | intraciliary transport involved in cilium assembly; intraciliary retrograde transport | intraciliary retrograde transport; microtubule-based movement | 0.135 |
| evgtrinLocDN32719c2g1t2 | intraciliary retrograde transport | intraciliary retrograde transport; microtubule-based movement | 0.199 |
| evgtrinLocDN28925c0g1t1 | intraciliary retrograde transport | intraciliary retrograde transport; microtubule-based movement | 0.131 |
| evgtrinLocDN31604c0g1t1 | microtubule-based movement | microtubule-based movement | 0.180 |
| evgtrinLocDN29990c0g1t2 | plus-end-directed vesicle transport along microtubule | microtubule-based movement | -0.140 |
| evgtrinLocDN37139c0g1t2 | microtubule-based movement | microtubule-based movement | 0.227 |
| evgtrinLocDN37139c0g2t1 | microtubule-based movement | microtubule-based movement | 0.151 |
| evgtrinLocDN32580c5g1t3 | microtubule-based movement | microtubule-based movement | -0.120 |
| evgtrinLocDN36762c6g2t2 | microtubule-based movement | microtubule-based movement | 0.191 |
| evgtrinLocDN38625c0g1t1 | chorion-containing eggshell formation | chorion-containing eggshell formation | -0.230 |
| evgtrinLocDN27146c8g2t1 | chorion-containing eggshell formation | chorion-containing eggshell formation | -0.146 |
| evgtrinLocDN29511c1g1t2 | copper ion homeostasis; cellular transition metal ion homeostasis | transition metal ion homeostasis | 0.149 |
| evgtrinLocDN36113c0g1t6 | iron ion homeostasis | transition metal ion homeostasis | -0.120 |
| evgtrinLocDN29153c4g1t3 | allantoin catabolic process | allantoin metabolic process; drug catabolic process | 0.184 |
| evgtrinLocDN30569c1g1t15 | allantoin catabolic process | allantoin metabolic process; drug catabolic process | 0.205 |
| evgtrinLocDN26098c0g1t1 | glycine decarboxylation via glycine cleavage system | drug catabolic process | 0.138 |
| evgtrinLocDN34135c2g2t2 | hydrogen peroxide catabolic process, response to oxidative stress | drug catabolic process | -0.126 |
| evgtrinLocDN31868c2g3t2 | chitin metabolic process | chitin metabolic process | 0.129 |
| evgtrinLocDN30862c9g1t7 | chitin metabolic process | chitin metabolic process | 0.141 |
| evgtrinLocDN38560c1g1t1 | chitin metabolic process | chitin metabolic process | -0.175 |
| evgtrinLocDN26516c0g3t2 | chitin metabolic process | chitin metabolic process | -0.196 |
| evgtrinLocDN25324c0g1t4 | chitin metabolic process | chitin metabolic process | 0.250 |
| evgtrinLocDN26665c13g2t1 | chitin metabolic process | chitin metabolic process | 0.297 |
| evgtrinLocDN33464c5g2t1 | chitin metabolic process | chitin metabolic process | 0.159 |
| evgtrinLocDN36412c0g1t18 | transposition, DNA-mediated | transposition, DNA-mediated | -0.123 |
| evgtrinLocDN35802c7g1t3 | transposition, DNA-mediated | transposition, DNA-mediated | -0.120 |
| evgtrinLocDN34439c0g2t3 | transposition, DNA-mediated | transposition, DNA-mediated | -0.136 |
| evgtrinLocDN36073c0g1t1 | transposition, DNA-mediated | transposition, DNA-mediated | 0.118 |
| evgtrinLocDN34933c0g1t1 | transposition, DNA-mediated | transposition, DNA-mediated | -0.171 |
| evgtrinLocDN36328c4g2t6 | transposition, DNA-mediated | transposition, DNA-mediated | -0.189 |

**Table S9.** Transcripts with annotated protein products within the percent of highest absolute loadings on RDA1.

| Transcript ID | Predicted protein product | Loading |
|---|---|---|
| evgtrinLocDN38595c4g1t1 | Transmembrane protein 135 | -0.425 |
| evgtrinLocDN29132c0g2t1 | Moderately methionine rich storage protein a | -0.376 |
| evgtrinLocDN31552c9g1t6 | Peripherin-2 like protein | -0.345 |
| evgtrinLocDN31767c0g1t2 | Moderately methionine rich storage protein a | -0.304 |
| evgtrinLocDN28466c0g2t3 | Lachesin | -0.272 |
| evgtrinLocDN27069c0g1t1 | Arylphorin subunit alpha | -0.233 |
| evgtrinLocDN34876c1g2t1 | Intraflagellar transport 52 homolog | 0.229 |
| evgtrinLocDN34466c4g4t2 | Gustatory receptor 22† | -0.229 |

† Pannzer2 predicted protein product corrected or refined after NCBI BLAST comparison.

**Table S10.** The GO biological process annotations of the transcripts associated with RDA4 that contribute to the enrichment of the biological processes in table S7.

| Transcript ID | GO BP annotation | GO BP enrichment contribution | Load. |
|---|---|---|---|
| evgtrinLocDN31645c0g1t1 | phosphatidylinositol-mediated signaling, phosphatidylinositol phosphorylation | phosphatidylinositol-mediated signaling, phosphatidylinositol phosphorylation | -0.086 |
| evgtrinLocDN36453c0g1t2 | phosphatidylinositol-mediated signaling, phosphatidylinositol phosphorylation | phosphatidylinositol-mediated signaling, phosphatidylinositol phosphorylation | -0.130 |
| evgtrinLocDN28658c0g1t2 | phosphatidylinositol-mediated signaling, phosphatidylinositol-3-phosphate biosynthetic process, phosphatidylinositol phosphorylation | phosphatidylinositol-mediated signaling, phosphatidylinositol phosphorylation, phosphatidylinositol-3-phosphate biosynthetic process | -0.054 |
| evgtrinLocDN25162c0g1t1 | phosphatidylinositol-mediated signaling, phosphatidylinositol phosphorylation | phosphatidylinositol-mediated signaling, phosphatidylinositol phosphorylation | -0.060 |
| evgtrinLocDN25042c0g1t2 | phosphatidylinositol-mediated signaling, phosphatidylinositol-3-phosphate biosynthetic process, phosphatidylinositol phosphorylation | phosphatidylinositol-mediated signaling, phosphatidylinositol phosphorylation, phosphatidylinositol-3-phosphate biosynthetic process | -0.063 |
| evgtrinLocDN23883c0g1t1 | protein localization to plasma membrane, phosphatidylinositol phosphorylation | phosphatidylinositol phosphorylation | -0.073 |
| evgtrinLocDN27172c5g1t2 | basolateral protein secretion, basement membrane constituent secretion, basal protein localization, regulation of basement membrane organization | basolateral protein secretion, regulation of basement membrane organization, basement membrane constituent secretion | -0.056 |
| evgtrinLocDN28461c0g1t1 | sensory organ precursor cell division, basolateral protein secretion, basement membrane constituent secretion, positive regulation of Notch signaling pathway | basolateral protein secretion, regulation of basement membrane organization, basement membrane constituent secretion | -0.062 |
| evgtrinLocDN29053c0g2t1 | basolateral protein secretion, basement membrane constituent secretion, basal protein localization, regulation of basement membrane organization, regulation of Rab protein signal transduction, basement membrane assembly, positive regulation of Golgi to plasma membrane protein transport | basolateral protein secretion, regulation of basement membrane organization, basement membrane constituent secretion, small GTPase mediated signal transduction | -0.076 |
| evgtrinLocDN34825c0g1t1 | small GTPase mediated signal transduction | small GTPase mediated signal transduction | -0.116 |
| evgtrinLocDN25616c0g1t1 | small GTPase mediated signal transduction | small GTPase mediated signal transduction | -0.082 |
| evgtrinLocDN36429c0g1t1 | regulation of ARF protein signal transduction | small GTPase mediated signal transduction | -0.085 |
| evgtrinLocDN29144c0g1t1 | small GTPase mediated signal transduction | small GTPase mediated signal transduction | -0.075 |

**Table S10 cont.**

| Transcript ID | GO BP annotation | GO BP enrichment contribution | Load. |
|---|---|---|---|
| evgtrinLocDN31236c0g1t1 | positive regulation of GTPase activity, regulation of small GTPase mediated signal transduction | small GTPase mediated signal transduction | -0.075 |
| evgtrinLocDN26095c0g3t1 | small GTPase mediated signal transduction | small GTPase mediated signal transduction | -0.072 |
| evgtrinLocDN30744c0g1t1 | small GTPase mediated signal transduction | small GTPase mediated signal transduction | -0.066 |
| evgtrinLocDN24761c0g1t1 | small GTPase mediated signal transduction | small GTPase mediated signal transduction | -0.073 |
| evgtrinLocDN32761c0g1t1 | small GTPase mediated signal transduction | small GTPase mediated signal transduction | -0.089 |
| evgtrinLocDN24985c0g1t2 | regulation of small GTPase mediated signal transduction, positive regulation of GTPase activity | small GTPase mediated signal transduction | -0.059 |
| evgtrinLocDN32450c0g1t1 | calcium ion transmembrane transport | calcium ion transmembrane transport | -0.060 |
| evgtrinLocDN25281c0g1t1 | calcium ion transmembrane transport | calcium ion transmembrane transport | -0.058 |
| evgtrinLocDN38307c0g1t4 | inositol phosphate-mediated signaling, calcium ion transmembrane transport | calcium ion transmembrane transport | -0.096 |
| evgtrinLocDN36351c0g1t2 | release of sequestered calcium ion into cytosol | calcium ion transmembrane transport | -0.097 |
| evgtrinLocDN25889c0g1t2 | regulation of ion transmembrane transport, calcium ion transmembrane transport | calcium ion transmembrane transport, regulation of ion transmembrane transport | -0.057 |
| evgtrinLocDN31628c0g1t2 5 | calcium ion transmembrane transport, regulation of ion transmembrane transport | calcium ion transmembrane transport, regulation of ion transmembrane transport | -0.066 |
| evgtrinLocDN38414c0g1t8 | regulation of ion transmembrane transport, calcium ion transmembrane transport | calcium ion transmembrane transport, regulation of ion transmembrane transport | 0.076 |
| evgtrinLocDN33995c1g1t4 | regulation of ion transmembrane transport, potassium ion transmembrane transport | regulation of ion transmembrane transport | 0.053 |
| evgtrinLocDN24552c0g1t2 | potassium ion transmembrane transport, regulation of ion transmembrane transport | regulation of ion transmembrane transport | -0.056 |
| evgtrinLocDN29306c7g1t1 | vacuolar acidification | regulation of intracellular pH | -0.080 |
| evgtrinLocDN36225c1g1t2 | phagosome reneutralization | regulation of intracellular pH | -0.084 |
| evgtrinLocDN30184c0g1t1 | vacuolar acidification, lumen formation, open tracheal system | regulation of intracellular pH | -0.096 |
| evgtrinLocDN37380c0g1t3 | protein retention in Golgi apparatus, protein targeting to vacuole | maintenance of protein location in cell | -0.153 |
| evgtrinLocDN37351c1g1t3 | cytoskeletal anchoring at plasma membrane, cell adhesion | maintenance of protein location in cell | -0.055 |
| evgtrinLocDN37216c0g1t1 | cytoskeletal anchoring at plasma membrane | maintenance of protein location in cell | -0.125 |
| evgtrinLocDN24238c0g1t1 | protein retention in Golgi apparatus, late endosome to vacuole transport | maintenance of protein location in cell | -0.064 |

**Table S11.** Transcripts with annotated protein products within the percent of highest absolute loadings on RDA4.

| Transcript ID | Predicted protein product | Loading |
|---|---|---|
| evgtrinLocDN31767c0g1t2 | Moderately methionine rich storage protein a | 0.160 |
| evgtrinLocDN33636c0g1t1 | HEAT repeat-containing protein 7A | -0.137 |
| evgtrinLocDN29132c0g2t1 | Moderately methionine rich storage protein a | 0.136 |
| evgtrinLocDN37744c0g1t5 | Transient receptor potential cation channel trpm | -0.120 |
| evgtrinLocDN35873c0g1t2 | Myosin-1A† | -0.119 |
| evgtrinLocDN34513c1g1t5 | Neurexin-1-alpha | -0.116 |
| evgtrinLocDN34142c4g1t3 | Moderately methionine rich storage protein a | 0.115 |
| evgtrinLocDN38360c1g1t1 | Serine/threonine-protein kinase TOR-like | -0.112 |
| evgtrinLocDN25410c0g1t1 | larval cuticle protein F1-like† | -0.110 |
| evgtrinLocDN22597c0g1t1 | Lysine-specific demethylase lid† | -0.107 |
| evgtrinLocDN27698c1g1t1 | Headcase protein | -0.105 |
| evgtrinLocDN38307c0g1t4 | Inositol 1,4,5-trisphosphate receptor | -0.096 |

† Pannzer2 predicted protein product corrected or refined after NCBI BLAST comparison.

**Table S12.** Statistically significantly enriched GO terms in transcripts differentially expressed within each family when exposed to host water limitation during early development.

| Family | GO ID | Term | $N_{annotated}$ | $N_{significant}$ | $P_{Fisher.elim}$ |
|---|---|---|---|---|---|
| F-1 | GO:0006030 | chitin metabolic process | 55 | 15 | <0.001 |
| | GO:0046033 | AMP metabolic process | 5 | 4 | <0.001 |
| | GO:0006275 | regulation of DNA replication | 5 | 3 | 0.002 |
| | GO:0010498 | proteasomal protein catabolic process | 35 | 7 | 0.006 |
| | GO:0006072 | glycerol-3-phosphate metabolic process | 7 | 3 | 0.008 |
| | GO:0007005 | mitochondrion organization | 47 | 8 | 0.009 |
| F-2 | GO:0034968 | histone lysine methylation | 31 | 8 | <0.001 |
| | GO:0007264 | small GTPase mediated signal transduction | 71 | 11 | 0.001 |
| | GO:0048015 | phosphatidylinositol-mediated signalling | 13 | 4 | 0.004 |
| | GO:0007424 | open tracheal system development | 14 | 4 | 0.005 |
| | GO:0051453 | regulation of intracellular pH | 8 | 3 | 0.007 |
| | GO:0031146 | SCF-dependent proteasomal ubiquitin-dependent prot. catab... | 8 | 3 | 0.007 |
| | GO:0065007 | biological regulation | 1096 | 85 | 0.007 |
| | GO:0046854 | phosphatidylinositol phosphorylation | 24 | 5 | 0.007 |
| | GO:1903508 | positive regulation of nucleic acid-templated transcription | 24 | 5 | 0.007 |
| | GO:0016579 | protein deubiquitination | 45 | 7 | 0.009 |
| F-5 | GO:0045087 | innate immune response | 14 | 4 | <0.001 |
| | GO:0009253 | peptidoglycan catabolic process | 12 | 3 | <0.001 |
| | GO:1901264 | carbohydrate derivative transport | 8 | 2 | 0.002 |
| F-6 | GO:0007264 | small GTPase mediated signal transduction | 71 | 24 | <0.001 |
| | GO:0048015 | phosphatidylinositol-mediated signalling | 13 | 8 | <0.001 |
| | GO:0046854 | phosphatidylinositol phosphorylation | 24 | 10 | 0.001 |
| | GO:0051603 | proteolysis involved in cellular protein... | 111 | 32 | 0.002 |
| | GO:0032869 | cellular response to insulin stimulus | 5 | 4 | 0.002 |
| | GO:0006511 | ubiquitin-dependent protein catabolic pr... | 97 | 25 | 0.003 |
| | GO:0043547 | positive regulation of GTPase activity | 58 | 17 | 0.003 |
| | GO:0016579 | protein deubiquitination | 45 | 14 | 0.004 |
| | GO:0032507 | maintenance of protein location in cell | 12 | 6 | 0.004 |
| | GO:0071711 | basement membrane organization | 6 | 4 | 0.005 |
| | GO:0016570 | histone modification | 87 | 22 | 0.006 |
| | GO:0043087 | regulation of GTPase activity | 72 | 23 | 0.010 |

**Table S13.** Information on the reliability of the SNP loci used for sex determination of the larvae.

| **Locus ID** | **Mean genot. score** | **Percent genot. failures** | **Heteroz. males** | **Heteroz. females** | **Repeatability success** | | **Previously found heteroz. females** | **Used** |
|---|---|---|---|---|---|---|---|---|
| | | | | | Sequenom duplicates | Sequenom vs. KASP | | |
| Titin1_R60_7999 | 0.935 | 14.1 | 0/6 | 1/6 | 9/10 | 12/12 | | Not |
| TRII_F37_S_8014 | 0.906 | 13.3 | 1/6 | 0/6 | 12/12 | 10/11 | | Not |
| Pro_192Y_8003 | 0.942 | 11.7 | 3/6 | 0/6 | 12/12 | 11/11 | | |
| s2308_1932_8013 | 0.925 | 10.1 | 0/6 | 0/6 | 13/13 | 11/11 | Yes | Not |
| snp_koe532_7996 | 0.959 | 7.6 | 1/6 | 1/6 | 12/13 | 11/11 | | Not |
| s4085_2312_8000 | 0.955 | 7.3 | 0/6 | 0/6 | 13/13 | 11/11 | | |
| s201_52818_7995 | 0.966 | 6.0 | 3/6 | 0/6 | 14/14 | 11/11 | | |
| s31_204583_8008 | 0.946 | 6.0 | 4/6 | 0/5 | 12/12 | 11/11 | | |
| s1764_1570_8012 | 0.911 | 4.1 | 0/4 | 0/6 | 12/12 | 9/9 | | |
| s734_29234_8009 | 0.965 | 3.3 | 2/6 | 0/6 | 14/14 | 11/11 | | |
| s2873_4167_8017 | 0.918 | 2.7 | 4/6 | 0/6 | 14/14 | 12/12 | | |
| s2188_2563_8016 | 0.882 | 2.2 | 0/6 | 0/6 | 14/14 | 12/12 | Yes | Not |
| s2428_5353_8015 | 0.969 | 2.2 | 2/6 | 0/6 | 14/14 | 11/11 | | |
| FLN_F63_SN_8002 | 0.982 | 1.6 | 3/6 | 0/6 | 13/13 | 11/11 | Yes | Unrel. |
| s1819_9187_8007 | 0.974 | 1.6 | 3/6 | 0/6 | 13/13 | 11/11 | | |
| s3244_5005_8005 | 0.984 | 1.6 | 0/6 | 0/6 | 13/13 | 11/11 | | |
| Lap_270Y_7997 | 0.984 | 1.1 | 3/6 | 0/6 | 14/14 | 12/12 | | |
| s4145_1014_8006 | 0.982 | 1.1 | 1/6 | 0/6 | 14/14 | 12/12 | | |
| snp_koe303_7998 | 0.988 | 1.1 | 2/6 | 0/6 | 13/13 | 11/11 | | |
| s2359_7598_8004 | 0.982 | 0.8 | 0/6 | 1/6 | 14/14 | 11/12 | | Unrel. |
| s523_12581_8011 | 0.984 | 0.8 | 4/6 | 0/6 | 14/14 | 12/12 | | |
| Titin2_408_8010 | 0.984 | 0.8 | 1/6 | 0/6 | 14/14 | 11/11 | Yes | Unrel. |
| w_scaffold_8001† | 0.895 | 8.7 | 0/6 | 3/6 | 13/13 | 12/12 | | Unrel. |

† Locus is pseudo-autosomal and is expected to be heterozygous in females, but not in males.
