## Supplementary methods for "Alternative developmental and transcriptomic responses to host plant water limitation in a butterfly metapopulation"

### *Detailed $^1$H-NMR protocol and processing of spectra*

After freeze drying the samples, we ground the dried pool samples in dry ice using a sterile pestle and prepared the finely ground tissue powder following the protocol by Kim et al. (2010). The $^1$H-NMR spectra of the *P. lanceolata* pool samples was recorded at the Finnish Biological NMR Center (Institute of Biotechnology, University of Helsinki, http://www.biocenter.helsinki.fi/bi/) using a Bruker Avance III HD NMR spectrometer (Bruker BioSpin, Germany) operated at $^1$H frequency of 850.4 MHz equipped with a cryogenic probe head. The $^1$H-NMR spectra were aquired at a 25 °C temperature with a “zgpr” pulse sequence.

After obtaining the spectra, we corrected the baseline, used automatic peak detection, binned the observed peaks into 0.04 ppm bins for all pool samples and normalized them according to the TSP signal variation and dry mass of the sample using MNOVA v.10.0.2 software (Mestrelab research S.L., Spain). We further annotated the peaks within the binned intervals based on signals of pure compounds for Aucubin, Catalpol and Verbascoside or from published characteristic signals (Agudelo-Romero et al., 2014; Gogna et al., 2015; Kim et al., 2010).

### *Detailed larval rearing protocol*

After hatching, we divided the larvae of each clutch into groups of twenty individuals and assigned them into one of the four treatments. In the beginning, we first provided each petri dish with a single 2.25cm$^2$ piece of control or water limited (see methods) *P. lanceolata* leaf tissue corresponding to the treatment assigned to the plate. To enable an *ad libitum* provisioning of host plant while minimizing access to old leaf tissue, we adjusted the number of provided leaf pieces according to the number of larvae in each developmental stage (based on observations derived in a pilot experiment). First and second larval instars received ca. 0.11cm$^2$ leaf tissue per larva (i.e. a single 2.25cm$^2$ piece daily) and third larval instars received 0.24cm$^2$ per larva for the first three days, after which we increased the feeding to ca. 0.34cm$^2$ per larva (i.e. three 2.25cm$^2$ pieces daily). Once larvae started entering the fourth larval instar, we kept on providing three 2.25cm2 pieces daily, but because in most larval groups some larvae entered diapause already in the fourth larval instar (Figure S4, Table S1) and because larvae were sampled for RNA, the amount of plant tissue available for each feeding individual was greater, typically between 0.5-0.7cm$^2$. Additionally, we monitored daily the consumption of the leaf pieces and reduced or increased the number of provided pieces according to consumption. If needed, we removed the previous day’s leftover leaf tissue.

*RNA and DNA extraction protocol*

Once removed from storage in -80°C the larvae were kept frozen in dry ice, transferred to 2ml tubes with 500µl of TRIzol and immersed in liquid nitrogen. The frozen samples were then homogenized, another 500µl of TRIzol was added and the samples incubated at room temperature for five minutes. After incubation, 0.2ml of chloroform was added, the solution was mixed thoroughly, and incubated for another 2-3 minutes at room temperature. The different phases were separated by first centrifuging the samples at 12 000 x g at 4°C for 15 min and then transferring the aqueous phase containing RNA to a separate tube. The aqueous phase continued to RNA extraction and the DNA containing organic/interphase continued to DNA extraction.

RNA extraction continued with adding 600µl phenol:chloroform (1:1) to the aqueous phase. The solution was then shaken thoroughly and centrifuged at 12 000 x g at 4°C for 10 min. The resulting upper phase was transferred to a new tube and RNA was precipitated with isopropanol by adding 50µl of 3M sodium acetate and 550µl isopropanol. The solution was incubated in room temperature for 10 minutes and centrifuged at 12 000 x g at 4°C for 10 min. If pellet could not be seen clearly, the centrifugation was extended. After removal of the supernatant, the pellet was washed with 1ml of 75% ethanol and centrifuged at 7 000 x g at 4°C for 5 min. The wash was discarded and the ethanol washing step repeated. Once washed, the pellet was air dried, immersed in 50µl of nuclease free water, incubated in 55-60°C for 10-15min and stored in -70°C until library preparation and sequencing.

DNA extraction continued by adding 0.3ml of 100% ethanol to the lower organic/interphase extracted from the TRIzol and chloroform part of the extraction procedure and mixing the solution thoroughly. After incubating for 2-3 min. the solution was centrifuged at 2000 x g at 4°C for 5 min, the supernatant was discarded and the resulting DNA pellet was air dried for 5-10min. After this, the extraction continued using QIAamp DNA Mini Kit (Qiagen®) extraction kit. The pellet was immersed in 180µl of Buffer ATL, 20µl of proteinase K was added, and the solution incubated overnight at 56°C in a thermo shaker. After incubation, 100µl of nuclease free water and 4 µl of RNase A was added and the solution incubated at room temperature for 2min. After this 200µl of Buffer AL was added and incubation continued at 70°C for 10min, after which 200µl of 100% ethanol was added to the solution. The resulting mixture was added to a spin column (QIAamp Mini spin column) in a 2ml collection tube, centrifuged at 6000 x g for 1min. and the filtrate discarded. The spin column was subsequently washed twice with two different buffer solutions. First, 500µl Buffer AW1 was added into the spin column (in a clean 2ml tube), and centrifuged at 6000 x g for 1min. Then the filtrate was discarded, 500µl Buffer AW2 was added into the spin column, and the column (again in a clean 2ml tube) centrifuged first at 20 000 x g for 3min. and then (in a clean 2ml tube) at 20 000 x g for a minute. Finally, in the last step (repeated twice) the DNA was eluted from the column with 100 µl of Milli-Q purified water by first incubating the sample for 5min. at room temperature and then centrifuging the sample at 6000 x g for a minute.

*Sequenom sex determination*

In Lepidoptera, males are typically homogametic with respect to Z-chromosome and females typically have either a Z0, ZW or ZWW karyotype (Traut et al., 2008), and therefore only males are expected to be heterozygous in loci residing in the Z-chromosome. Using a Sequenom SNP panel, we sequenced a set of 22 Z-chromosome specific SNP loci for all collected individuals in families F-1, F-2, F-5 and F-6. Additionally, because in small populations with low levels of heterozygosity across the genome males can be homozygous in all Z-chromosome specific loci, we sequenced a single pseudo-autosomal locus (w_scaffold_8001), which is amplified from both Z and W chromosomes. In addition to estimating genotyping score and water controls in each genotyping plate, we used a four-fold reliability estimation of each SNP locus. First, to see if a locus can contribute to identifying the sex of the individual (i.e. is heterozygous only in males and homozygous in all females), we genotyped six known males and six known females that had been reared to adults in previous experiments. Second, as these individuals had previously been genotyped for the same loci using a different approach (KASP), we could estimate repeatability between different genotyping approaches. Third, we sequenced 14 samples twice to estimate repeatability of the sequencing. Fourth, we consulted an expert opinion from a lab technician who had designed the markers. Finally, we checked for previous experiences of reliability in terms of observing heterozygosity in the Z-chromosome associated markers in females.

We considered five loci unreliable to the extent that we decided to remove them from further analyses (Table S13). Locus *Titin1_R60_7999* failed in genotyping in over 14% of the cases and failed in reproducibility between different Sequenom duplicates. Similarly, locus *TRII_F37_S_8014* also failed in genotyping in several instances and failed in reproducibility between Sequenom and KASP duplicates. Locus *snp_ koe532_7996* was heterozygous in one known female and failed in reproducibility between different Sequenom duplicates. Loci *s2308_1932_8013* and *s2188_2563_8016* had been found to be heterozygous in several females and homozygous in most males during primer design, and thus we removed also those from any downstream analyses. Four additional loci also occasionally exhibited heterozygosity in Z-chromosome specific loci in our test females (*locus s2359_7598_8004*), or during primer design (*Titin2_408_8010* and *FLN_F63_SN_8002*). However, we decided not to exclude these markers as erroneously identifying a female as male only results in more conservative identification of females. We also decided to interpreted the single pseudo-autosomal locus (*w_scaffold_8001*) with caution as it frequently failed in genotyping.

We then sexed the individuals based on homozygosity across the 16 Z-chromosome specific markers and gave priority to individuals heterozygous in the single pseudo-autosomal locus. However, we decided not to categorically exclude individuals for which the pseudo-autosomal locus could not be genotyped or was homozygous, as it frequently failed in genotyping and was sometimes homozygous also in the test females (Table S13). After sexing, we checked that the families had roughly equal proportions of individuals identified as male and as female. The median numbers of heterozygous Z-choromosome specific loci in the individuals identified as the males were 2, 7, 5 and 7 for families F-1, F-2, F-5 and F-6, respectively.

*Sequence pre-processing*

After removal of Illumina adapter sequence and trimming of low quality sequence (Trimmomatic version 0.33, ) on average 96% (sd = 0.4%) of the reads were retained for each sample, leading to a minimum coverage of 12.8M reads per sample (mean = 16.7M, sd = 1.2M). We then proceeded to verify the family structure of the larvae, because individual first instar larvae can occasionally be accidentally displaced despite much care taken to maintain full-sib family structure in the studied families. We did this by determining pairwise genetic distances of the individuals from single nucleotide polymorphisms (SNPs) in the sequenced reads. To obtain the SNP genotypes of the individuals, we mapped the pre-processed sequence reads against the *M. cinxia* genome (Ahola et al., 2014) using STAR (Dobin et al., 2013), and called the SNPs using the software package SAMtools (version 1.8; Li et al., 2009). We then determined genetic distances and performed clustering based on neighbour-joining trees using the R packages "ape" (version 5.1; Paradis, Claude, & Strimmer,

2004) and “adegenet” (version 2.1.1; Jombart & Ahmed, 2011). As a result of this process a single individual from the first replicate larval group of constant drought treatment of family F-1 was identified as not belonging to any of the sequenced families and was removed from further analyses.

*Transcriptome build*

We used the full set of 78 sequenced individuals to build a *de novo* reference transcriptome. Prior transcriptome building, we trimmed the raw with trimmomatic (Bolger et al., 2014), and digitally normalised the samples using Trinity v2.6.5 (Grabherr et al., 2011). We then combined Trinity (v2.6.5) and Velvet / Oases (Schulz et al., 2012) to construct transcriptome assemblies. We ran Trinity with standard settings, but used a range of seven kmer sizes (21 to 71 bp) for Velvet / Oases, producing a separate assembly for each kmer size. We combined the resulting assemblies into a single fasta file, filtered the combined assembly using the EvidentialGene pipeline (Gilbert, 2013; http://arthropods.eugenes.org/EvidentialGene/trassembly.html), and removed contigs smaller than 200bp or expressed at levels less than a single normalized count per million.

*Transcript read counts*

We used RSEM with bowtie aligner specified for paired-end reads to quantify the expression levels for each of the transcripts in the *de novo* transcriptome. On average 67.8% of the reads for each sample mapped to the transcriptome (59.3% uniquely and 8.5% to multiple transcripts; Figure S8).

*References*


Agudelo-Romero, P., Ali, K., Choi, Y. H., Sousa, L., Verpoorte, R., Tiburcio, A. F., & Fortes, A. M. (2014). Perturbation of polyamine catabolism affects grape ripening of Vitis vinifera cv. Trincadeira. *Plant Physiology and Biochemistry*, *74*, 141–155. https://doi.org/10.1016/j.plaphy.2013.11.002

Ahola, V., Lehtonen, R., Somervuo, P., Salmela, L., Koskinen, P., Rastas, P., et al. (2014). The Glanville fritillary genome retains an ancient karyotype and reveals selective chromosomal fusions in Lepidoptera. *Nature Communications*, *5*(1), 4737. https://doi.org/10.1038/ncomms5737

Bolger, A. M., Lohse, M., & Usadel, B. (2014). Trimmomatic: a flexible trimmer for Illumina sequence data. *Bioinformatics*, *30*(15), 2114–2120. https://doi.org/10.1093/bioinformatics/btu170

Dobin, A., Davis, C. A., Schlesinger, F., Drenkow, J., Zaleski, C., Jha, S., et al. (2013). STAR: ultrafast universal RNA-seq aligner. *Bioinformatics*, *29*(1), 15–21. https://doi.org/10.1093/bioinformatics/bts635

Gilbert, D. (2013, June). EvidentialGene. *7th Annual Arthropod Genomics Symposium*. Notre Dame: Indiana University. https://doi.org/10.7490/F1000RESEARCH.1112594.1

Gogna, N., Hamid, N., & Dorai, K. (2015). Metabolomic profiling of the phytomedicinal constituents of *Carica papaya* L. leaves and seeds by 1H NMR spectroscopy and multivariate statistical analysis. *Journal of Pharmaceutical and Biomedical Analysis*, *115*, 74–85. https://doi.org/10.1016/j.jpba.2015.06.035

Grabherr, M. G., Haas, B. J., Yassour, M., Levin, J. Z., Thompson, D. A., Amit, I., et al. (2011). Full-length transcriptome assembly from RNA-Seq data without a reference genome. *Nature Biotechnology*, *29*(7), 644–652. https://doi.org/10.1038/nbt.1883

Jombart, T., & Ahmed, I. (2011). adegenet 1.3-1: new tools for the analysis of genome-wide SNP data. *Bioinformatics*, *27*(21), 3070–3071. https://doi.org/10.1093/bioinformatics/btr521

Kim, H. K., Choi, Y. H., & Verpoorte, R. (2010). NMR-based metabolomic analysis of plants. *Nature Protocols*, *5*(3), 536–549. https://doi.org/10.1038/nprot.2009.237

Li, H., Handsaker, B., Wysoker, A., Fennell, T., Ruan, J., Homer, N., et al. (2009). The sequence alignment/map format and SAMtools. *Bioinformatics*, *25*(16), 2078–2079. https://doi.org/10.1093/bioinformatics/btp352

Paradis, E., Claude, J., & Strimmer, K. (2004). APE: Analyses of phylogenetics and evolution in R language. *Bioinformatics*, *20*(2), 289–290. https://doi.org/10.1093/bioinformatics/btg412

Schulz, M. H., Zerbino, D. R., Vingron, M., & Birney, E. (2012). Oases: robust de novo RNA-seq assembly across the dynamic range of expression levels. *Bioinformatics*, *28*(8), 1086–1092. https://doi.org/10.1093/bioinformatics/bts094

Traut, W., Sahara, K., & Marec, F. (2008). Sex chromosomes and sex determination in Lepidoptera. *Sexual Development*, *1*(6), 332–346. https://doi.org/10.1159/000111765
